# Burden analysis of missense variants in 1,330 disease-associated genes on 3D provides insights into the mutation effects

**DOI:** 10.1101/693259

**Authors:** Sumaiya Iqbal, Jakob B. Jespersen, Eduardo Perez-Palma, Patrick May, David Hoksza, Henrike O. Heyne, Shehab S. Ahmed, Zaara T. Rifat, M. Sohel Rahman, Kasper Lage, Aarno Palotie, Jeffrey R. Cottrell, Florence F. Wagner, Mark J. Daly, Arthur J. Campbell, Dennis Lal

**Author notes:** these authors contributed equally to this work (Jakob B. Jespersen, Eduardo Perez-Palma, Patrick May).

## Abstract

Interpretation of the colossal number of genetic variants identified from sequencing applications is one of the major bottlenecks in clinical genetics, with the inference of the effect of amino acid-substituting missense variants on protein structure and function being especially challenging. Here we evaluated the burden of amino acids affected in pathogenic variants (n=32,923) compared to the variants (n=164,915) from the general population in 1,330 disease-associated genes on forty protein features using over 14,000 experimentally-solved 3D structures. By analyzing the whole gene/variant set jointly, we identified 18 features associated with 3D mutational hotspots that are generally important for protein fitness and stability. Individual analyses performed for twenty-four protein functional classes further revealed 240 characteristics of mutational hotspots in total, including new associations recapitulating the sheer diversity across proteins essential structural regions. We demonstrated that the function-specific features of variants correspond to the readouts of mutagenesis experiments and positively correlate with clinically-interpreted pathogenic and benign missense variants. Finally, we made our results available through a web server to foster accessibility and downstream research. Our findings represent a crucial step towards translational genetics, from highlighting the impact of mutations on protein structure to rationalizing the pathogenicity of variants in terms of the perturbed molecular mechanisms.

## Background

Recent advances in technologies have ushered in an era of rapid and cost-effective genome and exome sequencing. Genetic screening is now frequently applied in clinical practice, especially for the diagnosis of rare diseases and cancer, producing a plethora of benign and disease-associated genetic variants^1,2^. A large fraction of these clinically-derived variants has been captured in databases such as OMIM^3^, HGMD^4^, and ClinVar^5^. Simultaneously, millions of likely-benign or mildly disease-influencing variants from the general population have been cataloged and are accessible through the Exome Aggregation Consortium (ExAC)^6^ and in the genome Aggregation Database (gnomAD)^7^. Missense variation causes an amino acid substitution upon a single nucleotide variation in the protein-coding region of the genome, which can cause a Mendelian phenotype. However, missense variants can be either benign or disease-causing, and both types coexist in almost every human disease gene^6^. This underscores that functional interpretation of missense variants is particularly difficult, and some specific amino acid residues and their positions are essential for protein function and thus are vulnerable to substitutions. Exome sequencing studies can identify such protein-altering missense variants; however, the missing knowledge about the consequence of missense variants on protein structure and function severely limits the interpretation of clinical genetic screening, and our understanding of the implication of missense variants in disease phenotype.

Current algorithms for variant interpretation incorporate numerous annotations to assess the pathogenicity of missense variants, including the conservation of sequences through evolution^8–10^, human population variation information^11^, and protein structural information^12,13^. With the latest explosion of genetic variant data, machine learning algorithms can learn and predict variant pathogenicity with reasonable accuracy^14,15^. However, variant pathogenicity predictions, combined from multiple *in silico* tools, represent only one piece of supporting evidence (referred to as PP3^16^) in the variant interpretation guidelines as proposed by American College of Medical Genetics and Genomics (ACMG). This is primarily because of the similar underlying biases in these algorithms’ prediction outputs^16^, resulting from the use of similar variant sets. Further, the prediction scores do not provide biologically interpretable annotations that can be translated into possible variant-induced changes of underlying protein function.

Studying the effect of disease-associated missense variation on the protein structure and function represents a growing research area. The damaging consequence of amino acid substitution has been found to be associated with the structural property and localization of the amino acid on protein structure^17–19^. Cluster analysis of patient variants and structural conservation analysis have shown that functionally essential amino acids and regions are not randomly distributed^20–22^. In discrete studies, missense variants have been found to disrupt protein function by perturbing protein-protein interactions^23^, modifying functional residues^24^, destabilizing the entire protein fold^25,26^, or by substituting amino acid residues in protein cores^27,28^ and in protein-protein interfaces^29–31^. However, characterization of missense variants using a multiple line of evidence from protein structure, including physicochemical, functional, and 3D-structure based features, has not been performed using a common, large-scale dataset and single, unified pipeline. Such a thorough characterization of disease-causing and benign missense variants on 3D can serve as a valuable resource for translational genetics, providing many structural and functional insights into the impact of an amino acid substitution. Moreover, functionally essential protein regions and associated features are shared among proteins performing similar molecular function as well as can be diverse across different protein classes. And, there is no study available that provides a comparative overview of protein features associated with pathogenic and benign missense variants for different protein functional classes. Developing an inclusive annotation of structural, chemical, and functional features for missense variants for different protein classes thus can be a powerful resource for hypothesizing function-dependent variant pathogenicity.

In this study, we investigate the features of amino acid residues altered by missense variants on protein structure at scale. First, we built an automated pipeline to map variants from genomic location to protein 3D structure location. Using the framework, we mapped >500,000 missense variants in 5,850 genes onto >43,000 protein 3D structures from the protein data bank (PDB)^32^. The variant set was comprised of population missense variants from the gnomAD database^7^ and pathogenic missense variants from the ClinVar5 (pathogenic and likely-pathogenic) and HGMD^4^ (disease mutation) databases. Second, we annotated the amino acid positions of these genes with a set of seven protein features having forty feature subtypes. Third, we evaluated the burden of population and pathogenic missense variants in 1,330 disease-associated genes on these forty features. To do so relative to protein function, we carried out separate analyses with variants in genes grouped into 24 protein functional classes^33^. We then validated the effectiveness of identified pathogenic and population missense variant-associated features on independent variant sets, ascertained clinically and functionally.

## Results

### Description of dataset

In this study, we characterized missense variants with 3D coordinates in at least one protein structure using seven protein features of amino acid residues: (1) 3-class secondary structure, (2) 8-class secondary structure, (3) residue exposure level, (4) physicochemical property, (5) protein-protein interaction, (6) post-translational modification, and (7) UniProt-based functional features. These seven features further have forty subtypes (description of features in **Methods** and **Supplementary Note**). We performed the characterization jointly for all variants and independently for variants in groups of genes with similar molecular function. Initial collection of solved human protein structures from Protein Data Bank^32^ (accessed September 2017) resulted in 43,805 tertiary structures for 5,850 genes. For 5,724 genes, we found variants from the gnomAD database^6^ (hereafter referred to as population variants). For 1,466 and 1,673 genes, we obtained pathogenic and likely-pathogenic variants from ClinVar^6^ and disease mutations from HGMD^6^ databases, respectively (hereafter referred to as pathogenic variants). In total, we aggregated 1,485,579, 16,570 and 47,036 variants (unique amino acid alterations) from gnomAD^7^, ClinVar^5^ and HGMD^4^, respectively. Out of the available variants, 33% (count = 496,869), 49.1% (count = 8,137), and 65.3% (count = 30,730) variants from gnomAD, ClinVar, and HGMD, respectively, were mappable to at least one protein structure from PDB. Population variants were mappable for 4,987 and pathogenic variants for 1,330 human genes. We restricted the data set to 1,330 genes with both population and pathogenic variants to perform an unbiased comparative analysis. The final dataset was comprised of 164,915 population variants and 32,923 pathogenic missense variants across 1,330 genes (**Supplementary Table 1**), which were grouped into 24 protein classes (**Supplementary Table 2**, protein class ascertainment in **Methods**) to perform protein class-specific analysis. This missense variant and gene dataset are referred to as Disease-Associated Genes with Structure (DAGS1330) in this study. The workflow of this study is briefly presented in **Fig. 1** and the overview of the dataset is shown in **Table 1**.

**Fig. 1.**
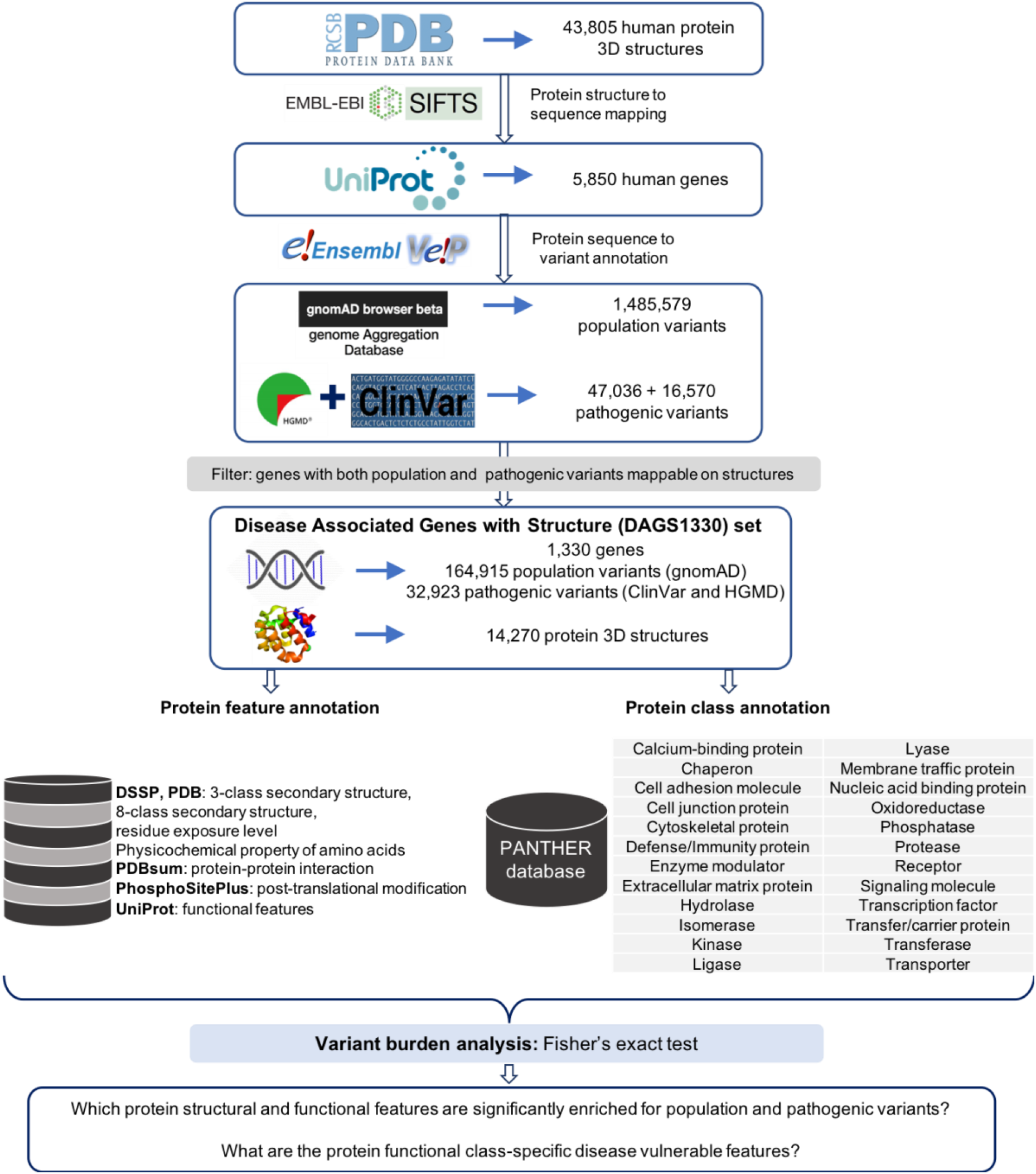
Flowchart of the study. Genetic variant to protein structure mapping, protein feature (from databases such as DSSP^34^, PDBsum^35^, PhosphoSitePlus^36^, UniProt^37^) and functional class (PANTHER^33^ database) annotations, and statistical analysis.

**Table 1.** Number of genes, missense variants (amino acid alterations), and protein structures in the dataset.

|  | Population variants | Pathogenic variants |  |
| --- | --- | --- | --- |
|  | gnomAD missense | ClinVar missense<br>(likely/-pathogenic) | HGMD missense<br>(disease mutation) |
| Genes with protein 3D structure | 5,724 | 1,466 | 1,673 |
| Number of missense variants in the genes<br>with protein 3D structure | 1,485,579 | 16,570 | 47,036 |
| Number of missense variants mapped on the<br>structure | 496,869 | 8,137 | 30,730 |
| Genes with missense variants mapped on the<br>structure | 4,897 | 1,330<sup>+,</sup> |  |
| Number of unique 3D structures on which the<br>variants were mapped | 29,870 | 14,270<sup>+</sup> |  |
| Number of missense variants in DAGS1330<br>genes that were mapped on the structure | 164,915<sup>+</sup> | 32,923<sup>+,</sup> |  |

<sup>+</sup> combined (unique) counts from HGMD and ClinVar databases.
<sup>+</sup> Genes and variants that are considered in the final statistical analysis, referred to as Disease Associated Genes with Structure (DAGS1330).

### Statistical analysis captures feature patterns of population and pathogenic missense variants on 3D structure across all protein classes

Our understanding of which protein features are more likely to be disease relevant is still in its infancy. To identify structural, physiochemical, and functional protein features that are associated with pathogenic and population missense variants, we performed burden analysis with seven features including in total 40 subtypes (ascertainment and definition of features in **Methods** and **Supplementary Note**) across DAGS1330 genes. Fisher’s Exact test was performed for individual feature subtypes of the population and pathogenic variants, quantifying the fold-enrichment and significance of the association. Notably, out of the seven features, physiochemical properties of amino acid and UniProt-based functional features could be curated from the protein sequence only. However, our dataset comprised of the missense variants that could be mapped on the protein 3D structure only, thus, the identified features depict the characteristics of variants on protein structure. All seven protein features showed a deviating distribution of population and pathogenic missense variants for at least one feature subtype. The direction of the association was heterogeneous across feature subtypes (**Fig. 2**). In total, 18 out of 40 (45%) protein feature subtypes were significantly enriched for pathogenic variants. The UniProt-based functional protein features had the highest number of feature subtypes (5/6) that were enriched for pathogenic variants (**Fig. 2**). In contrast, 14 out of 40 (35%) feature subtypes showed significant depletion of pathogenic variants and enrichment of population variants. The physiochemical properties showed enrichment of population variants in six out of eight subtypes. Eight protein feature subtypes (20%) did not show any specific association to pathogenic or population variants.

**Fig. 2.**
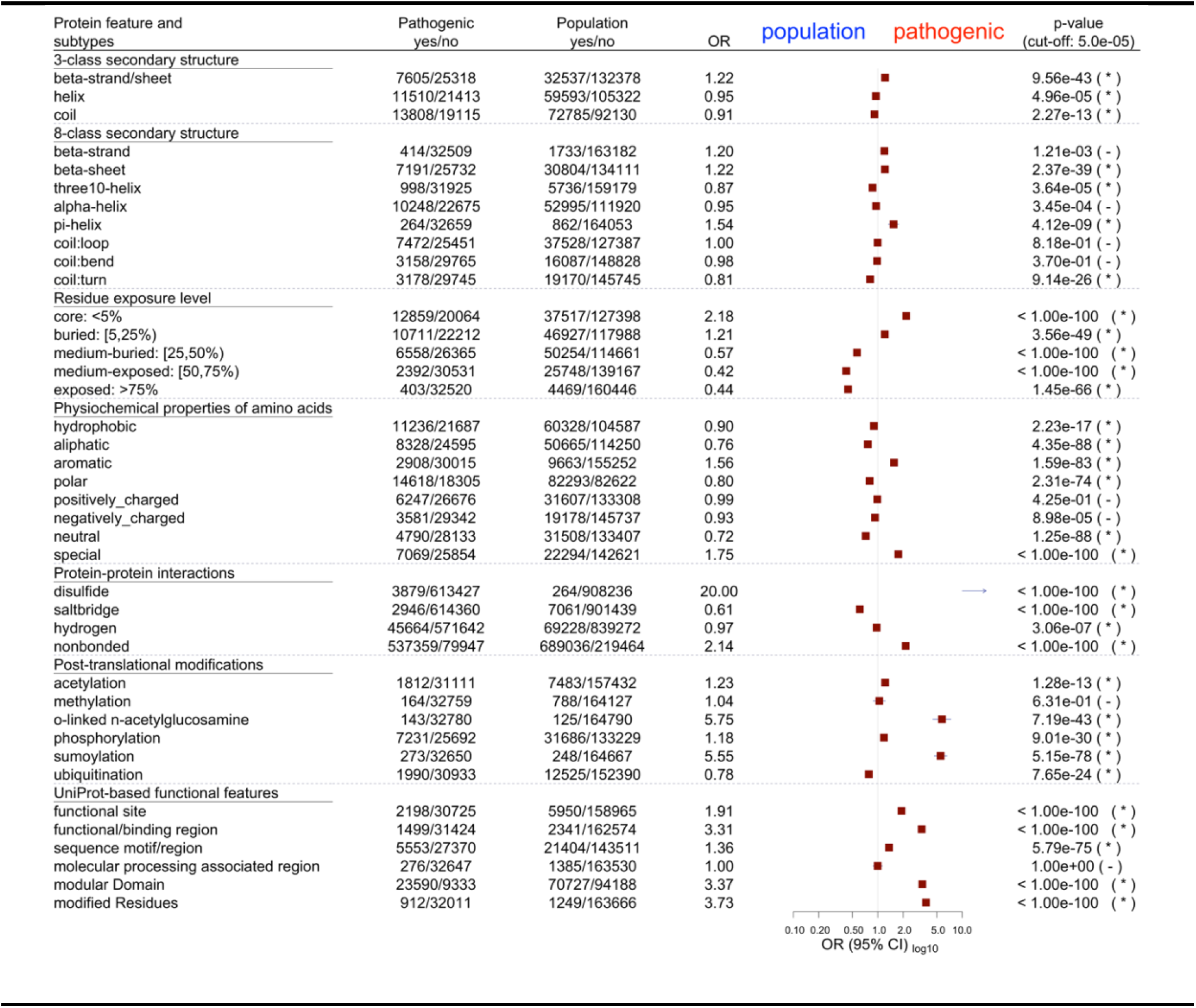
Burden of pathogenic and population variants in protein features, taking 32,923 pathogenic and 164,915 population variants in 1,330 genes. The plot shows results of the two-tailed Fisher’s Exact test for 40 feature subtypes. The y-axis outlines the feature names, followed by the feature counts with and without (yes/no) pathogenic and population variants, the odds ratio (OR) and the significance (p-value). The squares (brown) show the OR and the bars (blue) show the 95% confidence interval on the x-axis. OR=1 indicates the neutral value (no enrichment or depletion) while the OR > 1.0 (and < 1.0) indicates enrichment of pathogenic variants (and population variants). If the association is significant (p-value < pcut-off = 5.0e-05), the corresponding p-value is followed by (*), otherwise (-). For protein-protein interaction types, the feature counts correspond to all bond-annotations available for the amino acid residues with pathogenic and population variants. For the rest of the features, the counts correspond to the number of amino acid residues.

A wide range of effect sizes was observed for the enrichment of pathogenic variants across the 18 protein feature subtypes with odds ratio (OR) values ranging from 1.21 to 21.75 (**Fig. 2**). The top three features associated with pathogenic variants were intermolecular disulfide bonds (OR = 21.75), SUMOylation sites (OR = 5.75), and O.GlcNAc sites (OR = 5.55). We further investigated for a possible bias in our analysis due to the wide range of population to pathogenic variant ratio per gene (0.02 – 156.0). When the analyses were repeated using only the identical number of variants per gene (20,552 pathogenic and 20,552 population variants in 588 genes, details in **Supplementary Note**), the patterns of population and pathogenic missense-variant-associated features remained consistent for 35 out of 40 features, considering the direction of association (**Supplementary Fig. 1**). The major deviations compared to the full DAGS1330 dataset were the associations of salt-bridge ionic bond and hydrogen bond sites with pathogenic variants which were observed the opposite way for the full dataset (details in **Supplementary Note**).

### Protein class-specific analysis identifies function-dependent vulnerable features

Proteins are heterogenous in function and structure. We next explored if the features of pathogenic and population variants obtained from the full DAGS1330 dataset were consistent across different protein classes, having heterogenous molecular functions (see **Methods** for protein class ascertainment, **Supplementary Table 2** for protein class-specific genes and variant counts, **Supplementary Fig. 2 – 8** for protein class-specific Fisher’s Exact test outputs, and **Table 2** for the summary of the results). Through the protein functional class-specific analyses, we aim to capture both shared and unique function-specific features of pathogenic and population variants on protein structure.

**Table 2.**
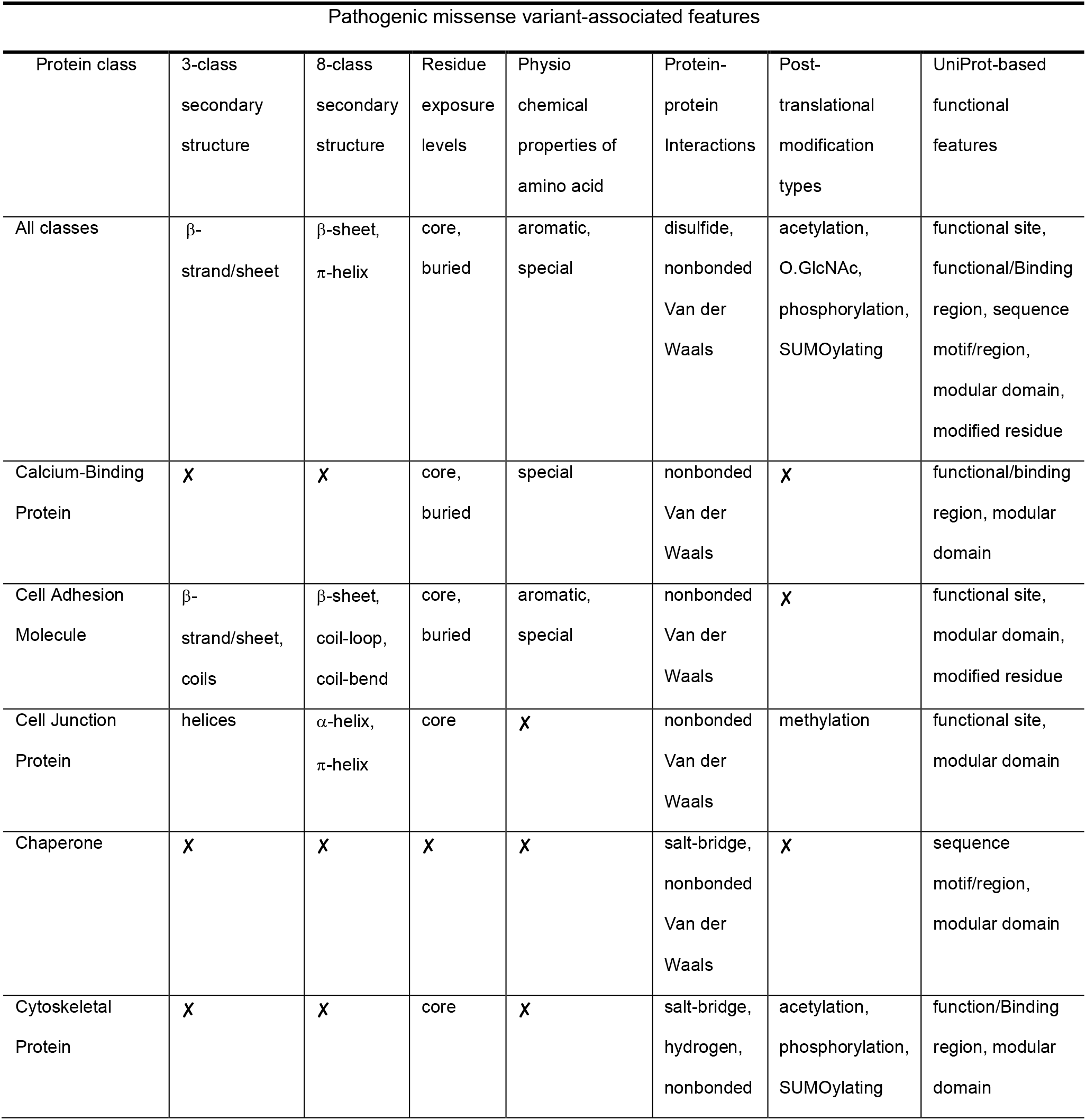
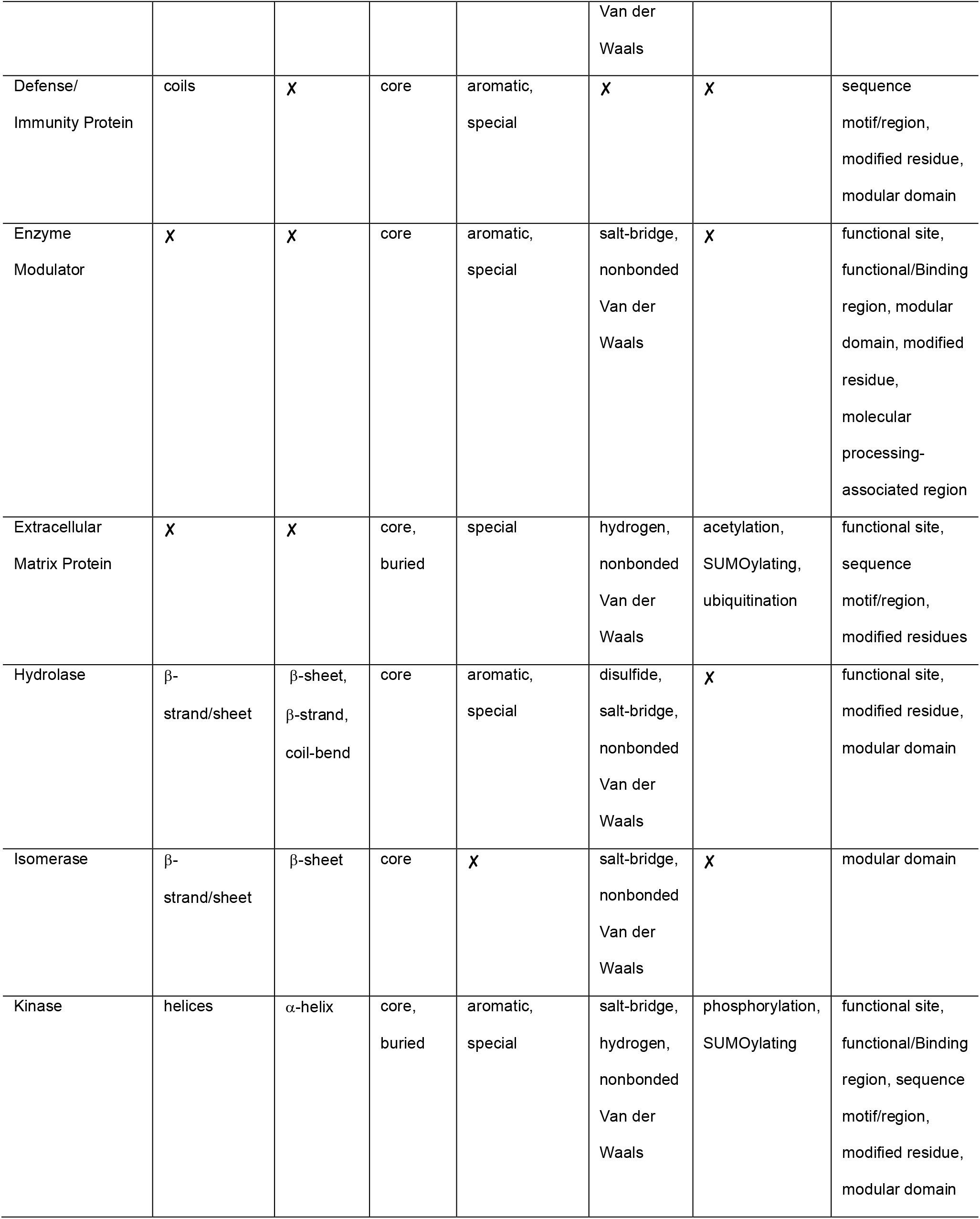
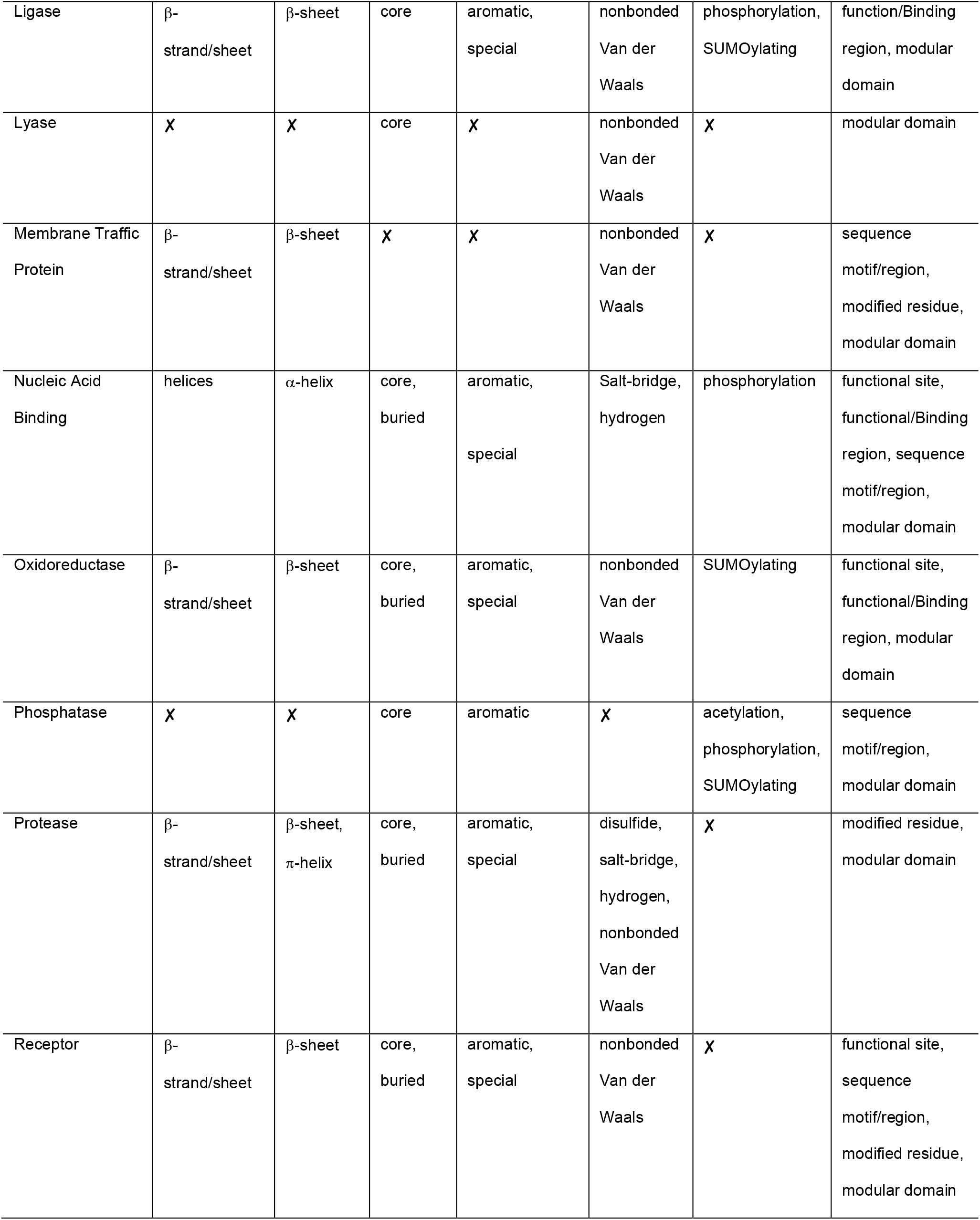
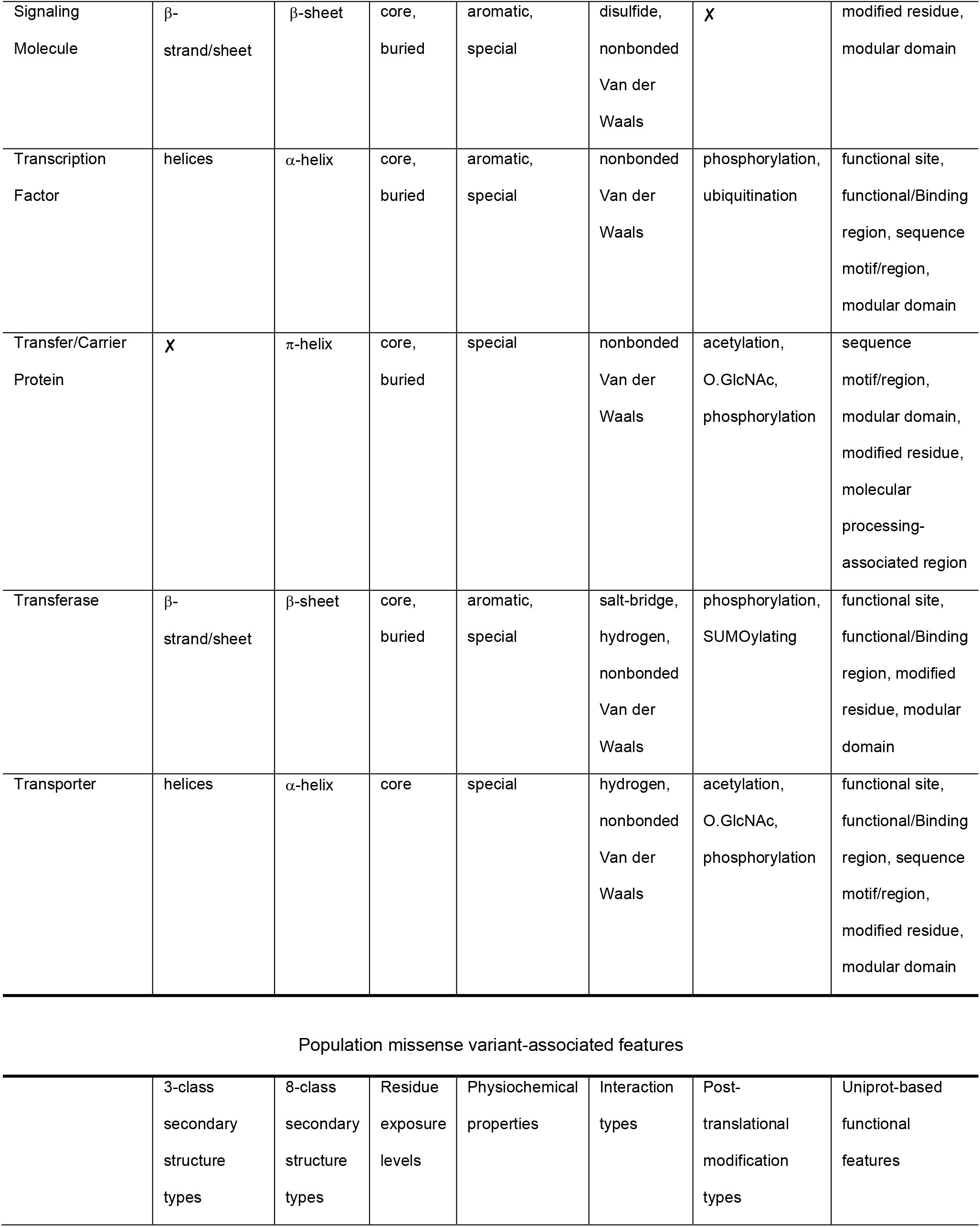
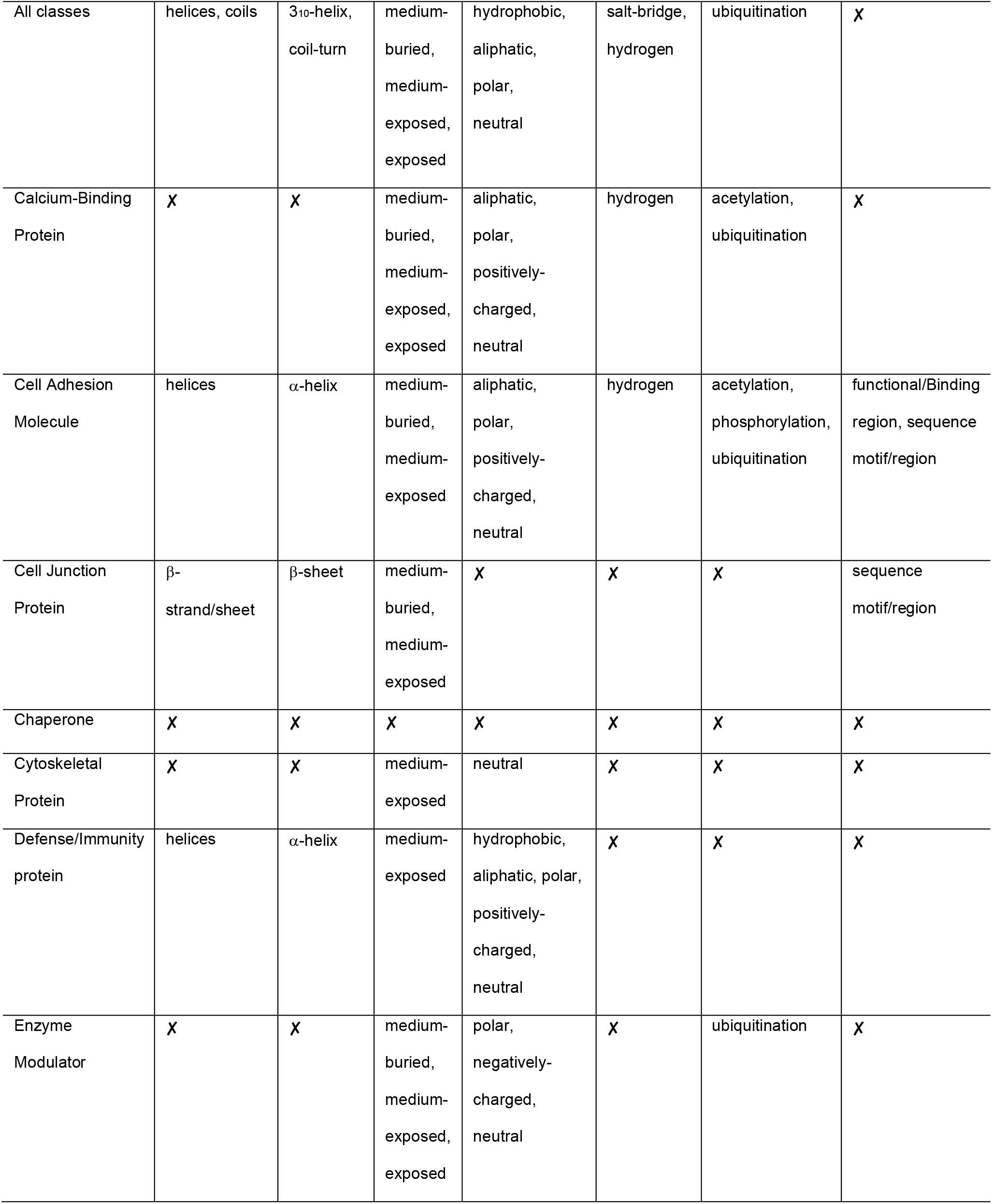
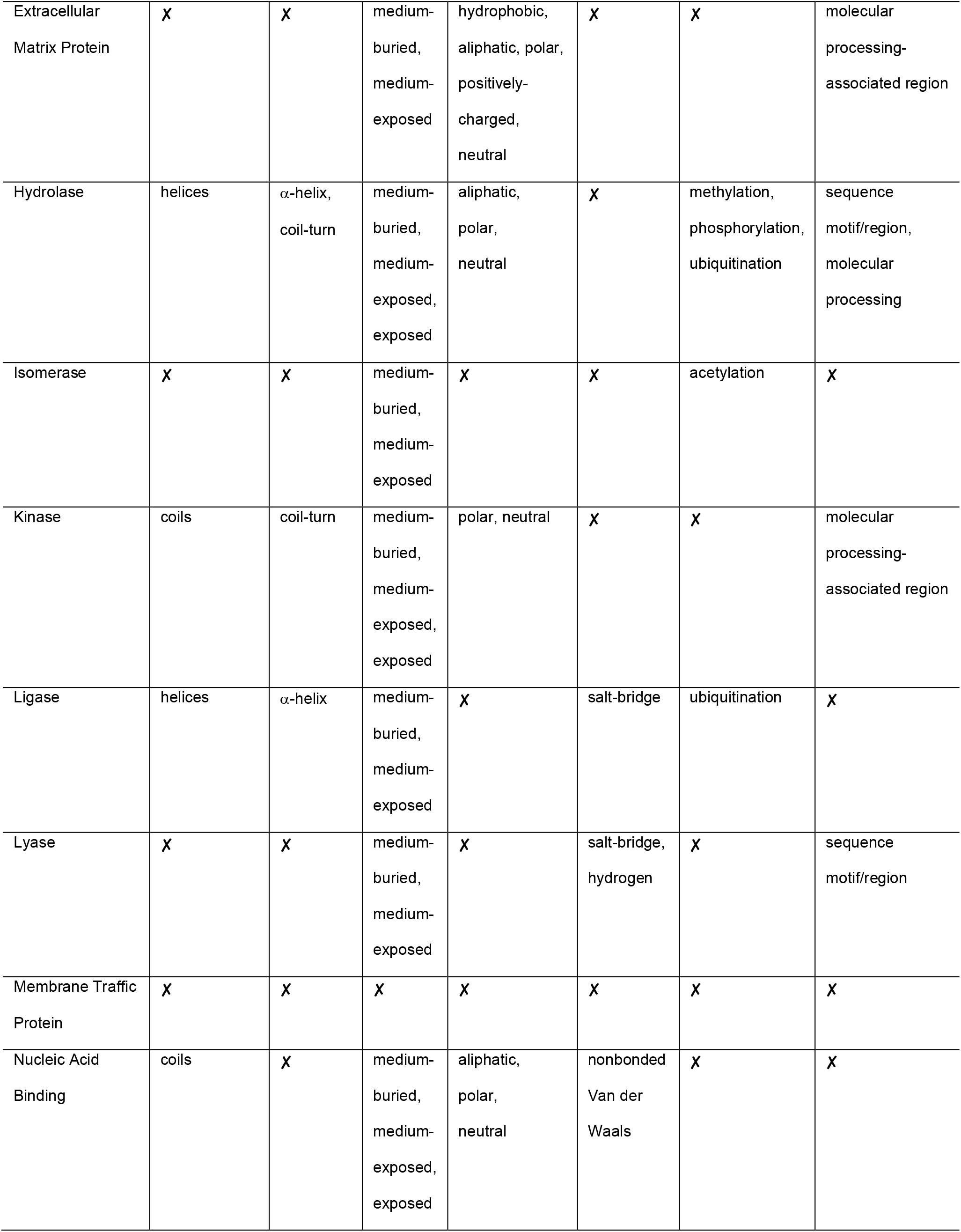
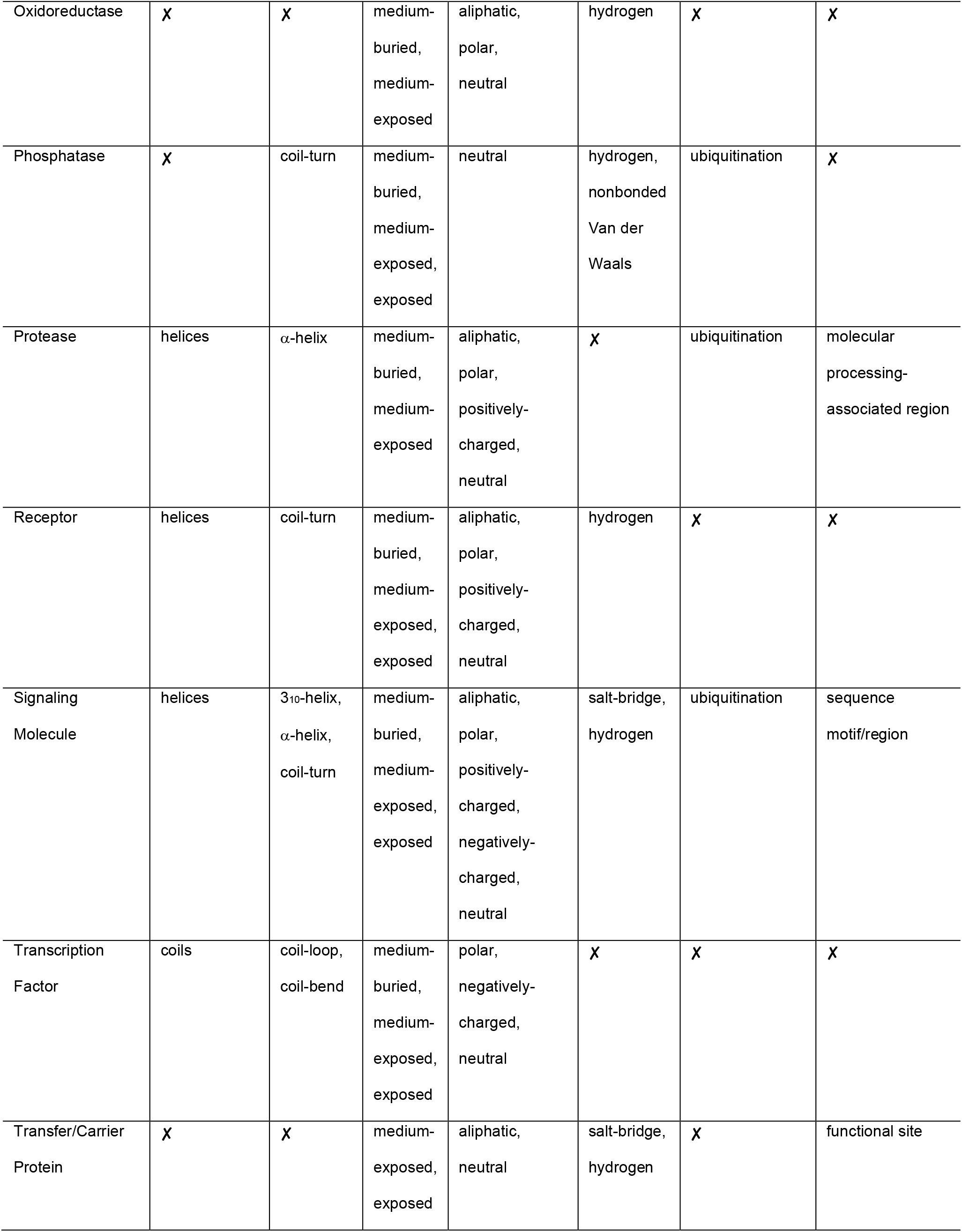
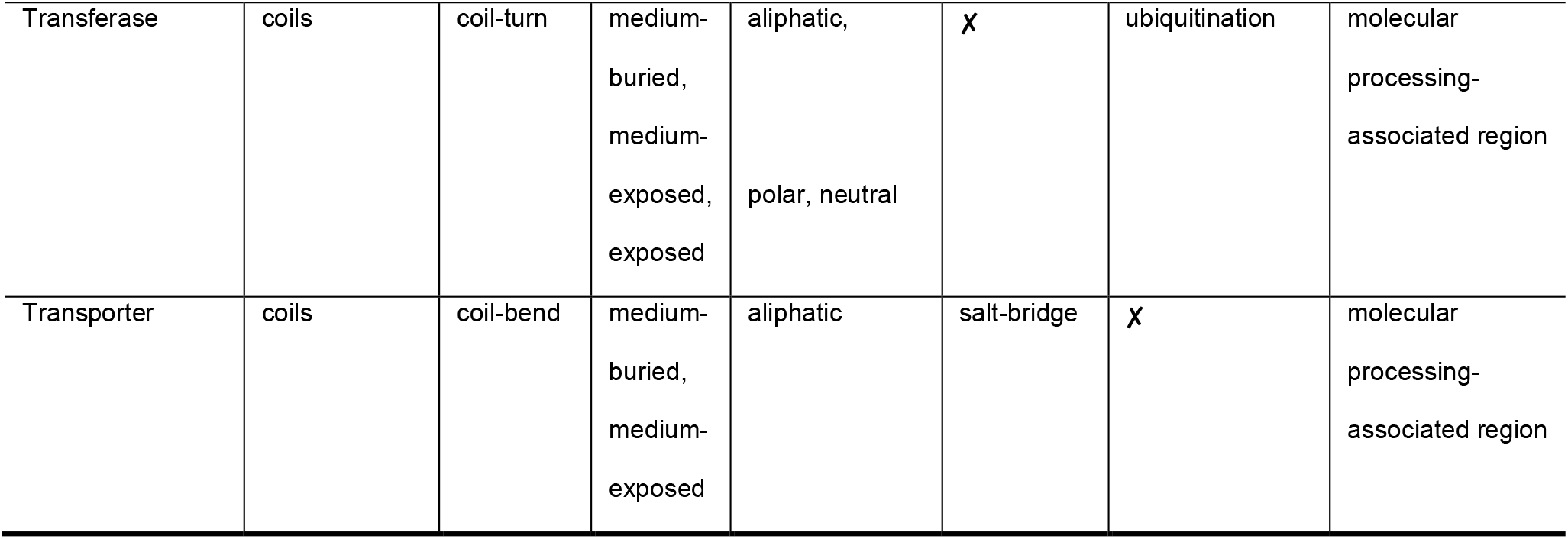
Protein class-specific pathogenic and population missense variant-associated features. Here, the columns are for seven different protein features where each cell outlines the feature subtypes that had significant burden of pathogenic and population variants (two-tailed Fisher’s Exact test, p < pcut-off, 5.0e-05). The ‘✘’ indicates that no subtype of the corresponding protein feature was significantly associated with a variant type.

We observed that pathogenic variants are enriched in β-strand/sheet residues for 20 out 24 of protein classes (**Supplementary Fig. 2a** and **3b**). This observation is consistent with the joint analysis with full DAGS1330 dataset (**Fig. 2**). In contrast, cell junction proteins displayed a deviating characteristic with enrichment of population variants in β-strand/sheet (OR = 0.37, p = 1.40e-13, **Supplementary Fig. 2a**), specifically, β-sheet (OR = 0.35, p = 3.91e-14, in **Supplementary Fig. 3b**). Moreover, pathogenic variants of cell adhesion molecules were significantly frequent in both stable β-sheet residues (OR = 1.72, p = 7.52e-25, **Supplementary Fig. 3b**) and flexible coils, specifically, random loops (OR = 1.28, p = 3.67e-06, **Supplementary Fig. 3f**), and bends (OR = 1.35, p = 2.19e-05, **Supplementary Fig. 3g**). The enrichment of pathogenic variants followed a negative gradient of exposure levels of amino acids (**Fig. 2**) and was consistent for the majority of the protein classes (**Supplementary Fig. 4**). Amino acid residues in the interior of the 3D structure with < 25% relative accessible surface area (RSA) were enriched with pathogenic variants (**Supplementary Fig. 4a-b**), whereas the residues with ≥ 25% RSA were enriched with population variants (**Supplementary Fig. 4c-e**). Among physiochemical property-based groups of amino acids, pathogenic variants were observed significantly more frequent on aromatic (tryptophan, tyrosine, phenylalanine) amino acids in 23 out of 24 protein classes, as well as on special amino acids (cysteine, proline, glycine) in all 24 protein classes (**Supplementary Fig. 5c** and **5h**). Specifically, the cell adhesion molecules, transcription factors, phosphatases, and signaling molecule proteins carried greater than 2-fold enrichments of pathogenic variants changing an aromatic amino acid. At positions of special amino acids, calcium-binding and extracellular matrix proteins showed enrichments of pathogenic variants with OR > 3.0 (p < 1.0e-100).

The inter-chain disulfide bond had the strongest enrichment in the joint analysis (OR = 21.75, p < 1.0e-100, **Fig. 2**). While such an association has been demonstrated 27, our protein class-specific analysis revealed that the enrichment was primarily directed by pathogenic variants from three protein classes alone (**Supplementary Fig. 6a**). In contrast, the non-bonded van der Waals interaction sites were enriched with pathogenic variants from all protein classes except phosphatases and nucleic acid binding proteins (**Supplementary Fig. 6d**). The pathogenic variants in kinases, nucleic acid binding and cytoskeletal proteins were found more frequent on salt-bridge and hydrogen bond sites (**Supplementary Fig. 6c-d**), which are protein class-specific features as these sites were found depleted of pathogenic variants for the full set (**Fig. 2**). Heterogeneous patterns of association between missense variants and post-translational modification (PTM) types were observed across different protein classes (**Supplementary Fig. 7**). The amino acid residues near phosphorylation sites were found to be enriched with pathogenic variants in joint analysis (**Fig. 2**); however, a differential pattern was observed for hydrolases (OR = 0.80, p = 7.29e-11) and cell adhesion molecules (OR = 0.58, p = 3.81e-15) with significant depletion of pathogenic variants around phosphorylation sites (**Supplementary Fig. 7d**). The cell junction proteins pathogenic variants showed an enrichment around methylation sites (OR = 9.10, p = 9.29e-11, **Supplementary Fig. 7b**), whereas there was no significant association observed for the full dataset. For nine out of twenty-four protein classes, we observed significant enrichment of population variants around ubiquitination sites; however, the trend was different for transcription factors (OR = 1.44, p = 1.77e-06) and extracellular matrix proteins (OR = 2.05, p = 2.51e-05), with an enrichment of pathogenic variants around ubiquitination sites (**Supplementary Fig. 7f**). Modular domains and modified residues were identified as shared functional features for a majority of the protein classes, twenty-three and twenty-two, respectively, out of twenty-four classes (**Supplementary Fig. 8**). The pathogenic variants of transporters, kinases, transcription factors, and nucleic acid binding proteins were observed to be associated with functional sites, binding regions and sequence motifs. The protein class-specific analysis also showed that only the transfer/carrier protein (OR = 8.90, p = 3.68e-09) and enzyme modulator (OR = 3.09, p = 6.20e-10) carry enrichment of pathogenic variants on protein regions involved in molecular processing and signaling (**Supplementary Fig. 8d**); however, we observed no association with this feature collectively for the full variant set (**Fig. 2**).

### Validation of 3D features associated to pathogenic and population missense variants on an independent set of ClinVar variants

Having characterized the pathogenic and population missense variants using 3D features (reported in **Table 2** and **Fig. 2**), we then carried out a comparison with an independent set of variants from ClinVar (see **Methods** for details) to validate the potential of our identified features for clinical interpretation of missense variants.

In order to quantify how deleterious an amino acid substitution is, we derived an index per residue based on the difference in the pathogenic and population variant-associated 3D features of the reference (altered) amino acid. The score is referred to as pathogenic 3D feature index (P3DFi) (see P3DFi computation in **Methods**). We expect that the amino acid residues located in vulnerable 3D sites will have a higher number of pathogenic variant-associated features (P3DFi > 0). Conversely, amino acids substituted in benign variants are expected to have a greater number of population variant-associated features (P3DFi < 0). Thus, we calculated the P3DFi values for amino acids affected by 18,094 pathogenic and 9,222 benign missense variants of 1,286 genes (see **Methods** for the preparation of validation dataset). We then binned the variants based on their P3DFi values (from less than −2 to greater than +2) and, as expected, the pathogenic and benign variants showed contrasting distributions across different P3DFi_DAGS1330_ values (**Fig. 3**). Overall, the pathogenic variants were 3.6-fold enriched (p < 1.0e-300) in positive P3DFi_DAGS1330_ values. Note that the most positive (P3DFi > 2) and negative (P3DFi < −2) score bins represent the 3D sites with highest and lowest difference between pathogenic- and population-associated 3D features (identified in this study). Remarkably, 82.7% (972 out of 1,175) of all variants in the highest ^**c**^;ore bin (P3DFi_DAGS1330_ > 2, **Fig. 3**) are pathogenic and 80.7% (1,048 out of 1,298) of all variants into the lowest score bin (P3DFi_DAGS1330_ < −2) are benign, leading to about 80% sensitivity and 84% specificity in the stratification of pathogenic and benign variants. Such a balanced sensitivity (true positive rate, pathogenic variants to possess positive P3DFi values) and specificity (true negative rate, benign variants to have negative P3DFi values) results in 82.7% precision and 17.2% false discovery rate in the variant classification.

**Fig. 3.**
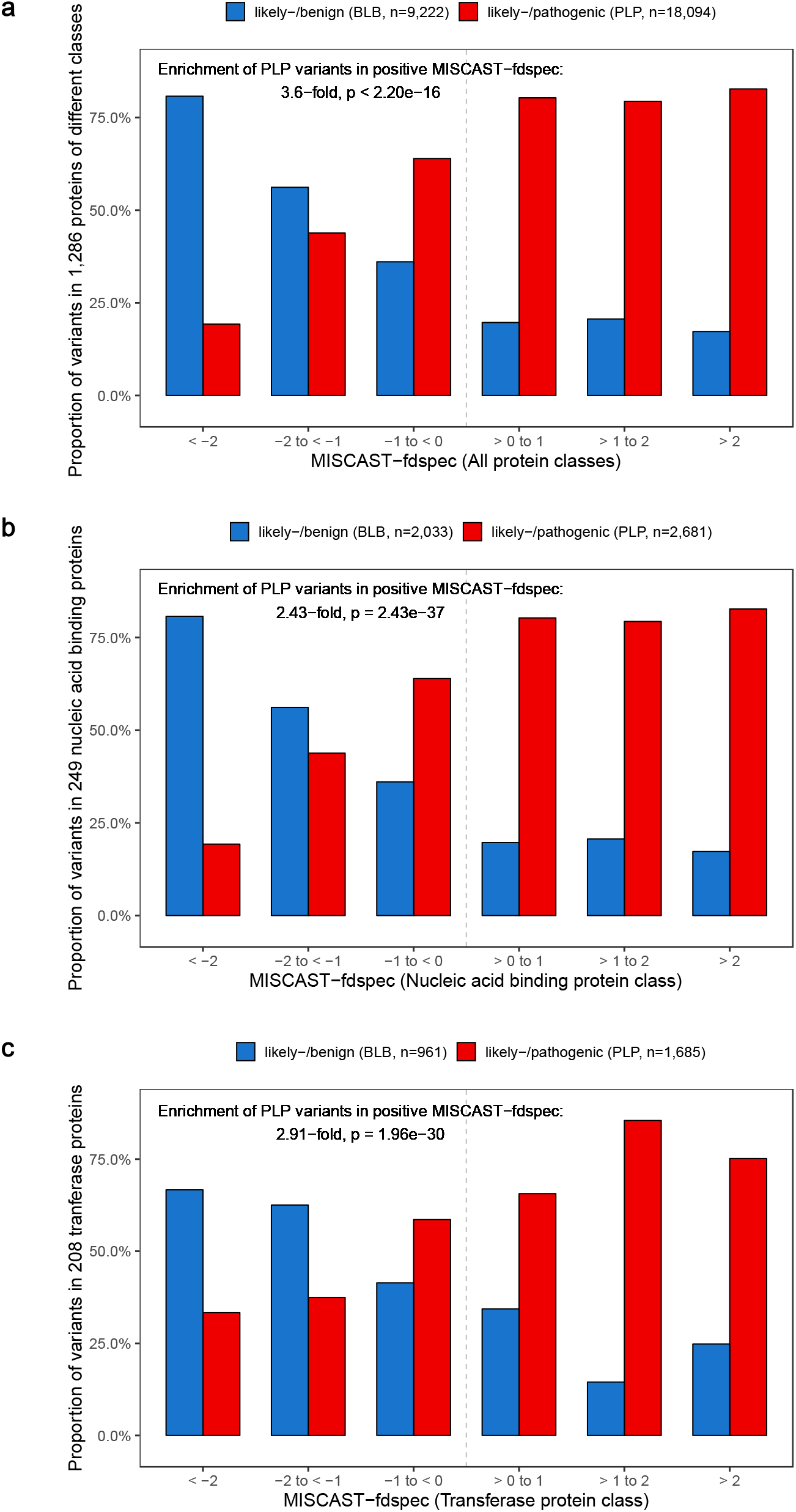
Validation of P3DFi values using clinically-ascertained independent variant set. **a,** Proportion of pathogenic and likely-pathogenic variants from ClinVar and disease mutations from HGMD in 1,286 genes (referred as PLP), independent from the training set, and benign and likely-benign variants from ClinVar (referred as BLB) in different P3DFi values, quantifying the difference between the number of pathogenic and population variant-associated features. The most confident positive P3DFi values (> 2) contains 4.9-times more PLP than BLB variants and that of confident negative P3DFi values (> 2) contains 4.2-times more BLB and PLP variants. **b,** Proportion of PLP and BLB variants in 249 nucleic acid binding genes in different P3DFi values, computed based on nucleic acid binding protein class-specific features. **c,** Proportion of PLP and BLB variants in 208 transferase genes in different P3DFi values bands, computed based on nucleic acid binding protein class-specific features.

Next, to provide example cases of similar evaluation of protein functional class-specific P3DFi (P3DFi_Protein class_) values, we selected two classes comprising the largest number of genes: nucleic acid-binding proteins (249 genes) and transferases (208 genes). Subsets of test variants of genes encoding for these two protein types were used in these evaluations, demonstrating a similar output as observed in **Fig. 3A**, i.e. the pathogenic and benign variants from both protein classes were more frequent in positive and negative P3DFi_Protein class_ values, respectively (results presented and discussed in **Fig. 3B-C**).

## Discussion

Here, we investigated forty different structural, physiochemical, and functional features of amino acids altered by missense variants in 1,330 genes (>80% annotated in OMIM) upon mapping of the variants onto 3D protein structures. We compared the burden of the population (gnomAD^7^) and pathogenic (combined ClinVar^5^ and HGMD^38^) variants in these forty features. As expected, pathogenic and population variants showed distinct distributions for many protein features. By joint analysis on all variants in 1,330 genes, we found eighteen pathogenic and fourteen population variant-associated protein features. Additionally, separate analyses on variants in genes of different protein classes resulted in novel function-specific features of pathogenic and population missense variants.

Currently a large number of disease-associated missense variants is available in publicly-accessible databases; however, a vast majority of these variants are of uncertain significance^5^, which fundamentally limits the clinical and research utility of genetic variation information. Given the complexity of variant classification, statistical approaches to objectively hypothesize the functional consequence of missense variants is pivotal^16^. Our analysis results will play a key role here as we compute the statistical burden of missense variants on experimentally solved protein structures, providing function-specific features and structural insights into the effect of missense variants at the molecular level, going beyond sequence- and conservation-based methods alone. It’s also worth noting that while the presence of variants in functional important protein domains has been widely utilized for variant interpretation^39,40^, we further consider critical structural and physicochemical features of the amino acid that may have functional implications on the protein structure and function upon perturbation. The current ACMG guidelines list the presence of variants in a mutational hot spot in less well-characterized protein regions or in a critical functional domain that is depleted in population variants as a moderate criterion (referred to as PM1)^16^ for variant pathogenicity assessment. However, domain information may not be the only protein-based feature that can be used as a determinant of variant pathogenicity. For reference, in our dataset, 28.3% of the pathogenic variants are not located in any protein domain according to UniProt annotation.

Our statistical analyses confirmed established association signatures between missense variants and protein features that have been found on a smaller scale and also illuminated novel protein class-specific signatures. For example, we observed an enrichment of pathogenic variants in β-strands/sheets conformation (**Fig. 2**) for all genes, thereby confirming the previous result that β-strands/sheets are more variant-intolerant and vulnerable to stabilization compared to helices upon sequence changes^41^. However, we found novel protein class-specific associations, such as pathogenic variants being significantly enriched on α-helices for five protein classes (**Supplementary Fig. 3d**). In cell junction proteins (OR = 10.54), transfer/carrier proteins (OR = 3.58), and proteases (OR = 2.71), pathogenic variants were more likely to mutate π-helix residues. The π-helical regions have been shown to undergo selection and often provide a functional advantage to proteins^42,43^; therefore, pathogenesis upon altering π-helices may be associated with perturbation of the protein functions. The protein core (Relative Solvent Area, RSA < 5%) is found to be a uniform descriptor of variant pathogenicity for all protein classes (**Supplementary Fig. 4a**), which is in line with established literature as well^27^. At the same time, for twenty-one out of twenty-four protein classes, distributions of RSA values for pathogenic missense variants showed higher maximum RSA values and accumulate a higher number of outliers (**Supplementary Fig. 10**). This result indicates the presence of pathogenic variants on the protein-protein surface^31^ that may perturb essential interaction sites by altering exposed residues.

On protein structure, the special amino acids cysteine (Cys, OR = 3.84, p < 1.00e-100) and glycine (Gly, OR = 1.76, p < 2.19e-154) were found to be predominantly mutated by pathogenic missense variation (**Supplementary Fig. 11**). The enrichment for Cys residues correlates well with the cogent association between variant pathogenicity and perturbation of covalent disulfide bond (a structure-based pathogenic variant-associated feature), as a Cys mutation is likely to impair such a bond^27^. However, there is a protein class-specific bias in the available disulfide bond annotations^35^ (**Supplementary Fig. 6a**). Specifically, 93.8% (3,642/3,879) of the disulfide bonds affected by a pathogenic variant were found in genes that encode for proteins transducing signals between cells. The other special amino acid, Gly, is the smallest amino acid having no side chain and is usually flexible. Greater than 67% of the Gly residues in our variant set is pliable coils and their substitutions by a larger amino acid is likely to have an impact on structure due to steric clashes^23^. All aromatic residues were found enriched for pathogenic variants (**Fig. 2**), with tryptophan (Trp) having the strongest association (OR = 3.27, p = 2.88e-126). Trp is the largest amino acid and it has also been shown that Trp mutations lead an increase in free energy of protein folding, making them more likely to destabilize the structure of protein^27^. Arginine (Arg) was also found to have a burden of pathogenic variants^31^ (**Supplementary Fig. 11**), while the other positively-charged amino acids (lysine and histidine) were found to be associated with population variants.

Pathogenic variants were, in general, found to spatially cluster near different post-translational modification (PTM) sites^44^; however, we identified several protein class-specific novel signatures. For example, the pathogenic variants in transcription factors, extracellular matrix proteins, and structural proteins were found to be enriched near ubiquitination sites (**Supplementary Fig. 8f**), which were found depleted of pathogenic variants by joint analysis (**Fig. 2**). Kinases alone carried 15% and 56% of the pathogenic variants near phosphorylation and SUMOylation sites. Expanding the six UniProt-based functional features into twenty-five constituent features (details in **Supplementary Note**), we identified twenty-one pathogenic variant-associated features. While some of these features, such as modular domains^45^, motifs, and binding sites^44^, are known to be associated with pathogenic variants, our study revealed new associations. A burden of pathogenic variants (OR = 3.9, p < 1.0e-100, **Supplementary Fig. 12**) on residues annotated by at least one functional feature versus no feature was also notable, showing putative association between variant pathogenicity and protein function.

It is important to note that we characterized the protein positions of missense variants that were mappable onto at least one protein 3D structure available in PDB^32^, which includes structures for only one-third of all human proteins. Many of these structures have partial coverage of the protein. Therefore, our study outcomes are not exclusive of the influence that may come from accounting for the structured part of the full protein sequence. Also, we could transfer a higher proportion of pathogenic variants (~60%) compared to the population variants (~33%) onto protein structure. Out of the 1,330 disease-associated genes (>80%, n = 1,077 annotated in OMIM), >55% of the genes had at least one structure with >50% of the sequence covered in the structure, and >75% of the ClinVar and HGMD pathogenic variants could be mapped onto the structure for those genes. This observation is consistent with the positive correlation observed between the number of known variants in proteins and the number of proteins being molecularly solved and reported in the literature^46^.

While performing the protein class-specific analyses, we also observed issues due to having smaller variant sets that could be mapped on the 3D structures. For one (lyase) and four (chaperons, cytoskeletal protein, defense/immunity proteins and membrane-traffic proteins) protein classes, we found three or fewer pathogenic and population variant-associated features (**Table 2**), respectively, after applying the stringent significance cut-off. In addition, the power of the P3DFi is affected by unavailability of 3D structure information of a reference amino acid, as, in that case, the quantitative score is computed based on the physicochemical and functional features only. In future work, we plan to include high-fidelity homology models and computational models of protein structures to scale up the generated resource to the full human proteome.

Currently, a variety of in silico tools aid in the interpretation of sequence variants, primarily providing a prediction score generated using different algorithms, training cohorts, and features^16^. The three most commonly used missense variant interpretation tools^16^ include PolyPhen2^8^, SIFT^10^ and Mutation Taster^47^. While these tools can stratify pathogenic variants from benign with a reasonable accuracy (65 – 80%)^48^, progress has not been made to provide users with biologically-relevant features of the location and context of missense variants within 3D structures. This is a problem in particular for molecular scientists who need a pre-selection of variants to study protein function. In this work, besides generating a quantitative score to rationalize missense variant pathogenicity, we also provide the relevant features to support the candidate variant selection by molecular biologists. Moreover, the gnomAD cohort has been leveraged for the first time in our work to assess genetic burden on protein features. Thus we believe that our results can serve as a powerful resource for the translation of personal genomics to precision medicine by translating genetic variant into 3D protein context: it can help to delineate variant pathogenicity, select candidate variants for functional assay, and aid in generating hypothesis for targeted drug development.

## Online Methods

### Protein structure and sequence collection

The Protein Data Bank (PDB)^32^ was mined on September 08, 2017 to collect all solved protein three-dimensional (3D) structures that were fully or partly (chimeric) indexed as homo sapiens. Our initial curation resulted in 43,805 protein structures, both single-chain and multiple-chain, including structures solved by six different experimental methods. Around 85% of the structures in this collection were solved by X-ray crystallography, of which over 50% of the structures in our collection are of resolution greater than 2.0 Å. The next highest proportion of the structures (~10%) were solved using nuclear magnetic resonance (NMR).

From UniProt^37^, the corresponding protein identifiers and gene name for the 43,805 protein structures were collected, which resulted in 5,870 unique protein identifiers and 5,850 unique gene names. Mapping of coordinates of amino acid residues in 3D structures to linear protein sequence was derived from the SIFTS database^49^.

### Missense variant collection and mapping on protein structure

Protein-coding missense variants in the general population were retrieved from the genome aggregation Database (gnomAD^6^) public release 2.0.2. Consolidated files for all available exomes and genomes (coding-only subset) were downloaded in Variant Call Format^50^ (VCFs) (http://gnomad.broadinstitute.org/downloads). Missense variant extraction was performed with vcftools based on the pre-annotated consequence “CSQ” field (VEP, Ensembl^51^ v92). All annotations refer to the human reference genome version GRCh37.p13/hg19. Entries passing gnomAD standard quality controls (Filter = “PASS” flag) and annotated to a canonical transcript were extracted (CSQ canonical = “YES” flag). The canonical transcript is defined as the longest CCDS translation with no stop codons according to Ensembl^51^. Exome and genome missense variants calls were consolidated into one single file using in-house Perl scripts (available upon request) matching identical genomic position and annotation.

We retrieved pathogenic missense variants from two sources: the ClinVar database (ClinVar)^5^, February, 2018 release and The Human Gene Mutation Database^38^ (HGMD^®^) Professional release 2017.2. ClinVar variants were downloaded directly from the ftp site (ftp://ftp.ncbi.nlm.nih.gov/pub/clinvar/) in table format. Missense variants were inferred through the analysis of the Human Genome Variation Society Sequence Variant Nomenclature field^45^ (HGVS, e.g. p.Gly1046Arg). To increase stringency, ClinVar variants were filtered to have only “Pathogenic” and/or “Likely Pathogenic” clinical consequence. Raw HGMD files were directly filtered for missense variants, high confidence calls (hgmd_confidence = “HIGH” flag), and disease-causing states (hgmd_variantType = “DM” flag). All annotations refer to the human reference genome version GRCh37.p13/hg19. Variants annotated to non-canonical transcripts were not considered.

Out of 5,850 genes with experimentally solved structures, we had gnomAD, ClinVar and HGMD missense variants for 5,724, 1,466 and 1,673 genes, respectively. Altogether, we collected 1,485,579; 16,570, and 47,036 amino acid alterations by missense variants from gnomAD, ClinVar and HGMD, respectively. We filtered out genes for which (i) canonical isoform protein sequences were not translated from canonical transcripts and (ii) amino acid residues affected by variants were not mappable to a protein structure. Thereafter, we had 496,869 gnomAD (33.4% of available variants), 8,137 ClinVar (49.1% of the available variants) and 30,730 HGMD (65.3% of the available variants) missense variants or amino acid alterations, respectively, that could be mapped at least one protein structure. Since ClinVar and HGMD are not mutually exclusive, we took the union of both resources, which was accomplished by comparison of their respective HGVS annotations using an in-house Perl script. Finally, we had 496,869 gnomAD (referred to as population/control) and 32,923 combined ClinVar and HGMD (referred to as pathogenic/case) missense amino acid variations from 4,897 and 1,330 genes, and these variants were mapped respectively onto 29,870 and 14,270 protein structures (PDB files). For comparative association analysis between population/pathogenic variants with protein features, we have used the 1,330 genes (Supplementary Table 1) only for which we had pathogenic (count = 32,923) and population (count = 164,915) variants. We refer to this dataset as the Disease-Associated Genes with Structure (DAGS1330). The full data overview is given in **Table 1**. The pipeline for mapping variants onto the protein 3D structures is shown in **Fig. 1**.

### Protein class annotation

The PANTHER (Protein ANalysis THrough Evolutionary Relationships) database^33^ was curated to classify genes by their molecular function, biological process, and protein class. The PANTHER Protein Class ontology was adapted from the PANTHER/X molecular function ontology.

We obtained the protein class annotations for 1,330 genes in the DAGS1330 dataset with population (gnomAD) and pathogenic (ClinVar and HGMD) variants mapped on the protein 3D structures. Initially, the genes were classified into 178 groups based on similar molecular function, 234 groups based on their involvement in similar biological processes, and into 206 different protein class groups. Finally, guided by the relationships defined among different protein classes (source: http://data.pantherdb.org/PANTHER13.1/ontology/Protein_Class_13.0), the genes were grouped into 24 major protein classes. After automatic annotation, we observed that 624 genes were not assigned to any specific protein class and so as to any protein major class. We then downloaded the Ensembl family description of all human genes (version 93) using BioMart (https://www.ensembl.org/biomart). After that, we manually annotated the 624 genes into those 24 major classes based on (i) the Ensemble family description, (ii) molecular function/biological process annotation previously collected from PANTHER or (iii) molecular function/biological process annotation available in UniProt as defined by the Gene Ontology consortium. **Supplementary Table 2** lists the 24 major classes and the number of genes, the number of protein 3D structures, and the count of population/pathogenic variants mapped on protein structures for each protein class. Note that each gene may not be assigned into a unique protein class group; therefore, the genes in each group are not mutually exclusive.

### Feature set mining and annotation

We annotated the amino acid residues of 1,330 genes with seven different features comprising of 40 feature subtypes. In the following, we introduce the features and we discuss how we curated these features and annotated the amino acid positions in the Supplementary Note.

1. 3-class secondary structure (subtype count: 3): (1) β-sheet/strand: β-strand and β-sheet; (2) Helices 3_10_-helix, α-helix, π-helix; (3) Coils - turn, bend, and random loops.
2. 8-class secondary structure (subtype count: 8): (1) β-strand; (2) β-sheet; (3) 3_10_-helix; (4) α-helix; (5) π-helix; (6) coil: turn; (7) coil: bend; (8) coil: loop.
3. Residue exposure level (subtype count: 5): (1) core (RSA < 5%); (2) buried (5% ≤ RSA < 25%); (3) medium-buried (25% ≤ RSA < 50%); (4) medium-exposed (50% ≤ RSA < 75%); (5) exposed (RSA ≥ 75%).
4. Physiochemical properties of amino acids (subtype count: 8): (1) aliphatic – alanine (Ala/A), isoleucine (Ile/I), leucine (Leu/L), methionine (Met/M), valine (Val/V); (2) aromatic – phenylalanine (Phe/F), tryptophan (Trp/W), tyrosine (Tyr/Y); (3) hydrophobic – aliphatic and aromatic amino acids; (4) positively-charged – arginine (Arg/R), histidine (His/H), lysine (Lys/K); (5) negatively-charged – aspartic acid (Asp/D), glutamic acid (Glu/E); (6) neutral – asparagine (Asn/N), glutamine (Gln/Q), serine (Ser/S), threonine (Thr/T); (7) polar – positively-charged, negatively-charged, and neutral amino acids; (8) special – proline (Pro/P), glycine (Gly/G), cystine (Cys/C).
5. Protein-protein interactions (subtype count: 4): (1) disulfide bond; (2) salt-bridge ionic interaction; (3) hydrogen bond; (4) nonbonded van der Waals interaction.
6. Post-translational modifications (subtype count: 6): (1) acetylation; (2) methylation; (3) O.GlcNAc, also known as β-linked N-acetylglucosamine; (4) phosphorylation; (5) SUMOylation; (6) ubiquitination.
7. UniProt-based functional features (subtype count: 6): (1) functional site – active site, metal binding site, binding site, site; (2) functional/binding region – zinc finger, DNA binding region, nucleotide phosphate binding region, calcium binding region; (3) sequence motif/region – region, repeat, coiled coil, motif; (4) modular domain – domain, topological domain, transmembrane, intramembrane; (5) molecular processing associated region – peptide, transit peptide, signal peptide, propeptide; (6) modified residue – modified residue, lipidation, disulfide bond, cross-link, glycosylation.

### Statistical analysis

For each of the 40 subtypes of seven protein features, we carried out the two-tailed Fisher Exact test on the 2×2 contingency matrix, populated with the total number of pathogenic and population variants of one specific protein feature subtype (helix within 3-class secondary structure) and the rest within a protein structure feature-group (‘not helix’ equals to ‘β-strand/sheet and coil’ within 3-class secondary structure). From this test, we obtained an estimate of fold enrichment, referred to as odds ratio (OR), along with the 95% confidence interval (CI) that shows whether the population/patient variants are relatively enriched or depleted in residues of any specific feature type on 3D structure. A value of OR equal to 1 indicates that there is no association between a specific type of variant with a particular feature, whereas a value of OR greater than 1 (or less than 1) indicates that the pathogenic variants are enriched (or depleted) in a particular type of feature subtype. Further, we calculated p-values showing the significance of association that were compared to the p_cut-off_. This analysis was also performed taking variants from all proteins and variants from the 24 different protein classes separately, and the output gives the features that are most likely to be associated to the pathogenic and population variants in general and also separately in different protein types. We defined pcut-off according to the Bonferroni correction for multiple testing in statistical analysis, thus pcut-off was set to 5.00e-05 (0.05/(40 * 25)) as we have 40 features on which we performed the association analysis taking all proteins and separately for 24 protein classes.

## Acknowledgements

We acknowledge Costin Leu, Giulio Genovese and Jon Bloom for insightful discussions that motivated some of the analysis presented in this manuscript. We are grateful to members of the Analytic and Translational Genetics Unit of Massachusetts General Hospital the medicinal chemistry group in the Stanley Center for useful comments and conversations. This work was supported by the Stanley Center for Psychiatric Research.

## Author Contributions

D.L., A.J.C. and F.F.W. conceived the research question. S.I., D.L., A.J.C. designed the study. S.I. led the study, performed data aggregation, statistical analyses and drafted the manuscript. S.I., J.B.J, E.P and P.M. collected the data. S.I., H.O.H., P.M., A.J.C., M.J.D. and D.L. contributed to analysis concepts and methods. S.I., P.M., A.J.C., M.J.D. and D.L. interpreted the results. S.I., D.H, E.P, S.S.A. and Z.T.R. developed the online resource. D.L., M.J.D., A.J.C. and P.M. supervised the study. D.L., M.J.D., A.J.C., P.M., K.L., M.S.R., H.O.H., A.P., F.F.W. and J.R.C. provided miscellaneous support to the research. All the authors revised and approved the final manuscript.

