## Supplementary Figures for "Burden analysis of missense variants in 1,330 disease-associated genes on 3D provides insights into the mutation effects"

### Insights into protein structural, physicochemical, and functional consequences of missense variants in 1,330 disease-associated human genes

Sumaiya Iqbal[1,2,3], Jakob B. Jespersen[4,5+], Eduardo Perez-Palma[6,+], Patrick May[7,+], David Hoksza[7,8], Henrike O. Heyne[1,2,3,9], Shehab S. Ahmed[10], Zaara T. Rifat[10], M. Sohel Rahman[10], Kasper Lage[1,4], Aarno Palotie[1,2,9], Jeffrey R. Cottrell[1], Florence F. Wagner[1,11], Mark J. Daly[1,2,3,9], Arthur J. Campbell[1,11,*], Dennis Lal[1,6,12,13,*]

[1]Stanley Center for Psychiatric Research, Broad Institute of MIT and Harvard, Cambridge, MA 02142, USA

[2]Program in Medical and Population Genetics, Broad Institute of MIT and Harvard, Cambridge, MA 02142, USA

[3]Analytic and Translational Genetics Unit, Massachusetts General Hospital, Boston, MA 02114, USA

[4]Department of Surgery, Massachusetts General Hospital, Boston, Massachusetts 02114, USA

[5]Department of Bio and Health Informatics, Technical University of Denmark, Lyngby, Denmark

[6]Cologne Center for Genomics, University of Cologne, Cologne, Germany

[7]Luxembourg Centre for Systems Biomedicine, University of Luxembourg, Esch-sur-Alzette, Luxembourg

[8]Department of Software Engineering, Faculty of Mathematics and Physics, Charles University, Prague, Czech Republic

[9]Institute for Molecular Medicine Finland (FIMM), University of Helsinki, 00100 Helsinki, Finland

[10]Computer Science and Engineering, Bangladesh University of Engineering and Technology, ECE Building, West Palashi, Dhaka-1205, Bangladesh

[11]Center for the Development of Therapeutics, Broad Institute of MIT and Harvard, Cambridge, MA 02142, USA

[12]Epilepsy Center, Neurological Institute, Cleveland Clinic, Cleveland, USA

[13]Genomic Medicine Institute, Lerner Research Institute Cleveland Clinic, US

[+]these authors contributed equally to this work

| Protein feature subtypes | Pathogenic yes/no | Population yes/no | OR | p-value (cut-off: 1.25e-03) |
|---|---|---|---|---|
| **3-class secondary structure** | | | | |
| beta-strand/sheet | 4672/15880 | 3848/16704 | 1.28 | 1.24e-23 ( * ) |
| helix | 7464/13088 | 7469/13083 | 1.00 | 9.67e-01 ( - ) |
| coil | 8416/12136 | 9235/11317 | 0.85 | 3.57e-16 ( * ) |
| **8-class secondary structure** | | | | |
| beta-strand | 271/20281 | 236/20316 | 1.15 | 1.29e-01 ( - ) |
| beta-sheet | 4401/16151 | 3612/16940 | 1.28 | 9.47e-23 ( * ) |
| three10-helix | 641/19911 | 673/19879 | 0.95 | 3.85e-01 ( - ) |
| alpha-helix | 6641/13911 | 6640/13912 | 1.00 | 1.00e+00 ( - ) |
| pi-helix | 182/20370 | 156/20396 | 1.17 | 1.72e-01 ( - ) |
| coil:loop | 4554/15998 | 4759/15793 | 0.94 | 1.62e-02 ( - ) |
| coil:bend | 1876/18676 | 2067/18485 | 0.90 | 1.46e-03 ( - ) |
| coil:turn | 1986/18566 | 2409/18143 | 0.81 | 1.58e-11 ( * ) |
| **Residue exposure levels** | | | | |
| core: <5% | 8101/12451 | 4424/16128 | 2.37 | < 1.00e-100 ( * ) |
| buried: [5,25%) | 6557/13995 | 5560/14992 | 1.26 | 4.32e-27 ( * ) |
| medium-buried: [25,50%) | 4114/16438 | 6482/14070 | 0.54 | < 1.00e-100 ( * ) |
| medium-exposed: [50,75%) | 1543/19009 | 3448/17104 | 0.40 | < 1.00e-100 ( * ) |
| exposed: >75% | 237/20315 | 638/19914 | 0.36 | 4.79e-44 ( * ) |
| **Physiochemical properties of amino acids** | | | | |
| hydrophobic | 6784/13768 | 7410/13142 | 0.87 | 8.91e-11 ( * ) |
| aliphatic | 5083/15469 | 6245/14307 | 0.75 | 1.19e-37 ( * ) |
| aromatic | 1701/18851 | 1165/19387 | 1.50 | 2.77e-25 ( * ) |
| polar | 9359/11193 | 10269/10283 | 0.84 | 2.77e-19 ( * ) |
| positively_charged | 4022/16530 | 3928/16624 | 1.03 | 2.45e-01 ( - ) |
| negatively_charged | 2277/18275 | 2405/18147 | 0.94 | 4.86e-02 ( - ) |
| neutral | 3060/17492 | 3936/16616 | 0.74 | 1.37e-30 ( * ) |
| special | 4409/16143 | 2873/17679 | 1.68 | 4.77e-88 ( * ) |
| **Protein-protein interactions** | | | | |
| disulfide | 3162/272487 | 220/163771 | 8.64 | < 1.00e-100 ( * ) |
| saltbridge | 2254/273395 | 1018/162973 | 1.32 | 1.08e-13 ( * ) |
| hydrogen | 22860/252789 | 12110/151881 | 1.13 | 2.97e-27 ( * ) |
| nonbonded | 230110/45539 | 133195/30796 | 1.17 | 7.89e-81 ( * ) |
| **Post-translational modifications** | | | | |
| acetylation | 991/19561 | 921/19631 | 1.08 | 1.06e-01 ( - ) |
| methylation | 125/20427 | 141/20411 | 0.89 | 3.56e-01 ( - ) |
| o-linked n-acetylglucosamine | 31/20521 | 41/20511 | 0.76 | 2.88e-01 ( - ) |
| phosphorylation | 4650/15902 | 4159/16393 | 1.15 | 3.84e-09 ( * ) |
| sumoylation | 89/20463 | 53/20499 | 1.68 | 3.12e-03 ( - ) |
| ubiquitination | 1282/19270 | 1231/19321 | 1.04 | 3.03e-01 ( - ) |
| **UniProt-based functional features** | | | | |
| functional site | 1363/19189 | 906/19646 | 1.54 | 5.08e-23 ( * ) |
| functional/binding region | 884/19668 | 288/20264 | 3.16 | 9.54e-73 ( * ) |
| sequence motif/region | 3243/17309 | 2080/18472 | 1.66 | 8.80e-66 ( * ) |
| molecular processing associated region | 207/20345 | 231/20321 | 0.90 | 2.69e-01 ( - ) |
| modular Domain | 13880/6672 | 10349/10203 | 2.05 | < 1.00e-100 ( * ) |
| modified Residues | 524/20028 | 183/20369 | 2.91 | 1.41e-39 ( * ) |

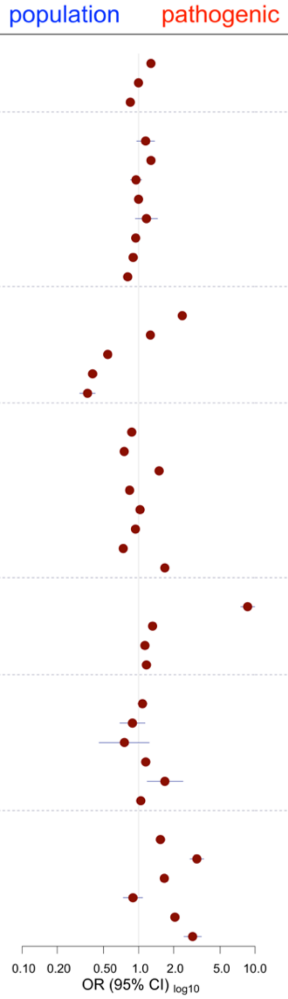


**Supplementary Fig. 1 | Burden of pathogenic and population variants in protein features, taking 20,552 pathogenic and 20,552 population variants in 588 genes.** The plot shows output of two-tailed Fisher's Exact test for 40 feature subtypes. The y-axis outlines the feature names, followed by the feature counts with and without (yes/no) pathogenic and population variants, the odds ratio (OR) and the significance (p-value). The squares (brown) show the OR and the bars (blue) show the 95% confidence interval on the x-axis. OR=1 is the neutral value of 1 (no enrichment or depletion) while the OR > 1.0 (and < 1.0) indicates enrichment of pathogenic variants (and population variants). If the association is significant (p-value < $p_{cut\text{-}off}$ = 1.25.0e-03), the corresponding p-value is followed by (*), otherwise (-). For protein-protein interaction types, the feature counts correspond to all bond-annotations available for the amino acid residues with pathogenic and population variants. For the rest of the features, the counts correspond to the number of amino acid residues.

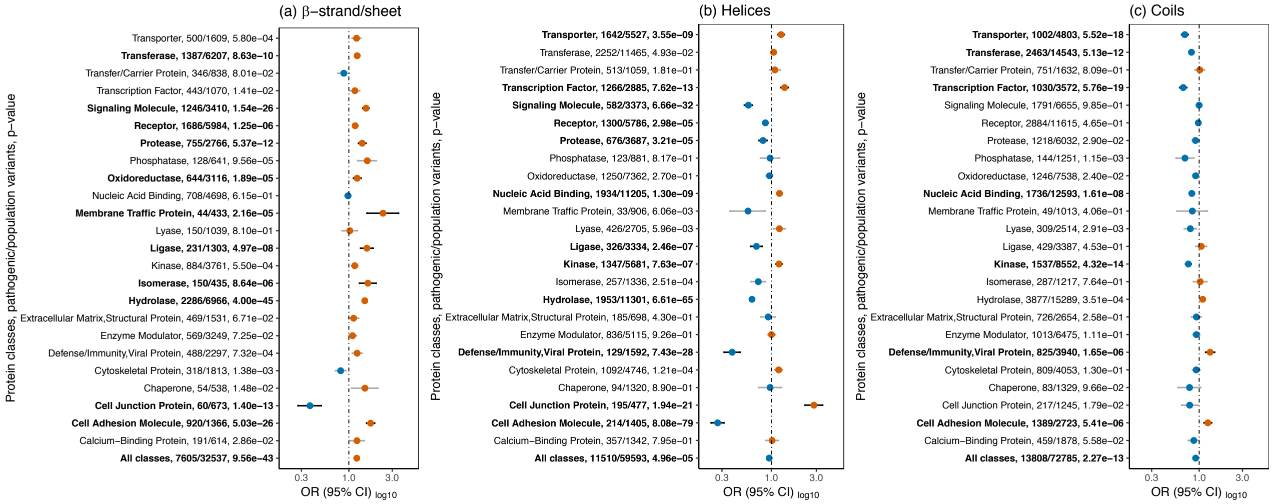


**Supplementary Fig. 2 | Protein-class-specific burden of pathogenic and population missense variants in the 3-class secondary structure feature.** In each plot, the odds ratio (OR) equal to 1 indicates there is no association (Fisher's Exact test) between a variant type and a feature subtype, whereas OR > 1 (orange) and OR < 1 (blue) indicates that the pathogenic and population variants, respectively, are enriched in a particular feature subtype. The minimum and maximum values of OR set to 0.0 and 20.0, respectively, for visualization. The x-axis labels (protein class, feature counts with pathogenic and population variants, p-values) and the horizontal bars showing the confidence intervals of OR are bold-faced for significant p-values, lower than the cut-off (5.50e-05) after the Bonferroni correction.

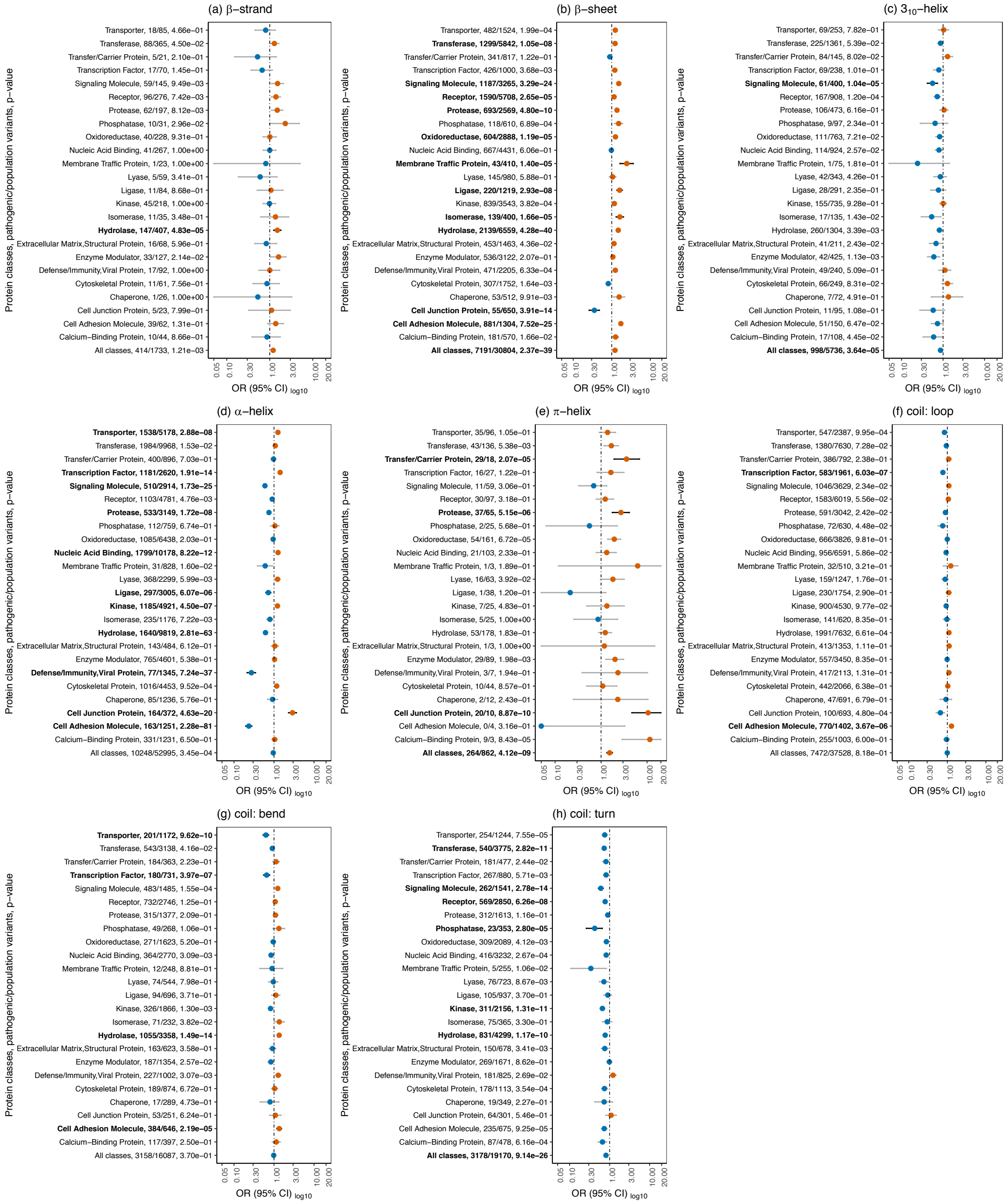


**Supplementary Fig. 3 | Protein-class-specific burden of pathogenic and population missense variants in the 8-class secondary structure feature.** In each plot, the odds ratio (OR) equal to 1 indicates there is no association (Fisher's Exact test) between a variant type and a feature subtype, whereas OR > 1 (orange) and OR < 1 (blue) indicates that the pathogenic and population variants, respectively, are enriched in a particular feature subtype. The minimum and maximum values of OR set to 0.0 and 20.0, respectively, for visualization. The x-axis labels (protein class, feature counts with pathogenic and population variants, p-values) and the horizontal bars showing the confidence intervals of OR are bold-faced for significant p-values, lower than the cut-off (5.50e-05) after the Bonferroni correction.

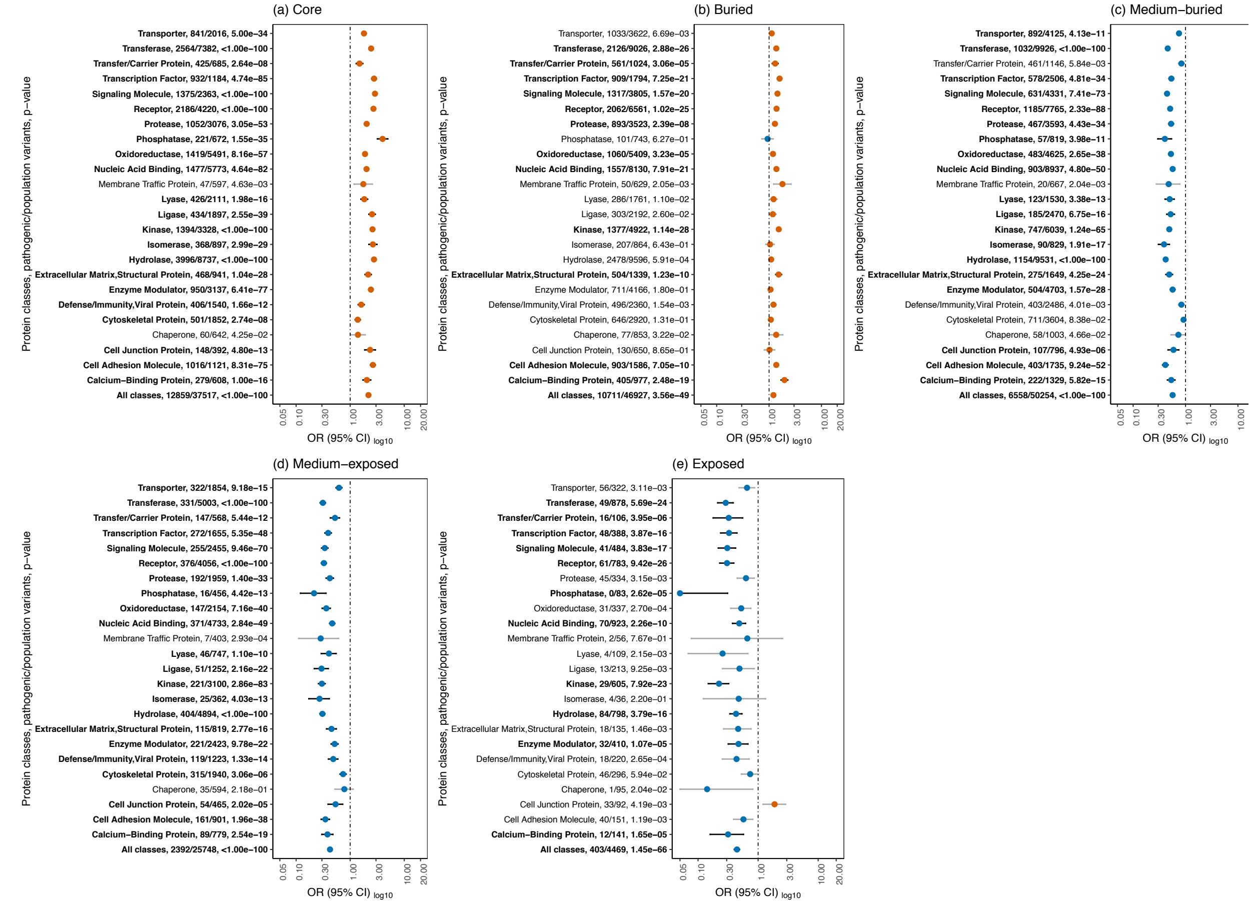


**Supplementary Fig. 4 | Protein-class-specific burden of pathogenic and population missense variants in the residue exposure level feature.** In each plot, the odds ratio (OR) equal to 1 indicates there is no association (Fisher's Exact test) between a variant type and a feature subtype, whereas OR > 1 (orange) and OR < 1 (blue) indicates that the pathogenic and population variants, respectively, are enriched in a particular feature subtype. The minimum and maximum values of OR set to 0.0 and 20.0, respectively, for visualization. The x-axis labels (protein class, feature counts with pathogenic and population variants, p-values) and the horizontal bars showing the confidence intervals of OR are bold-faced for significant p-values, lower than the cut-off (5.50e-05) after the Bonferroni correction.

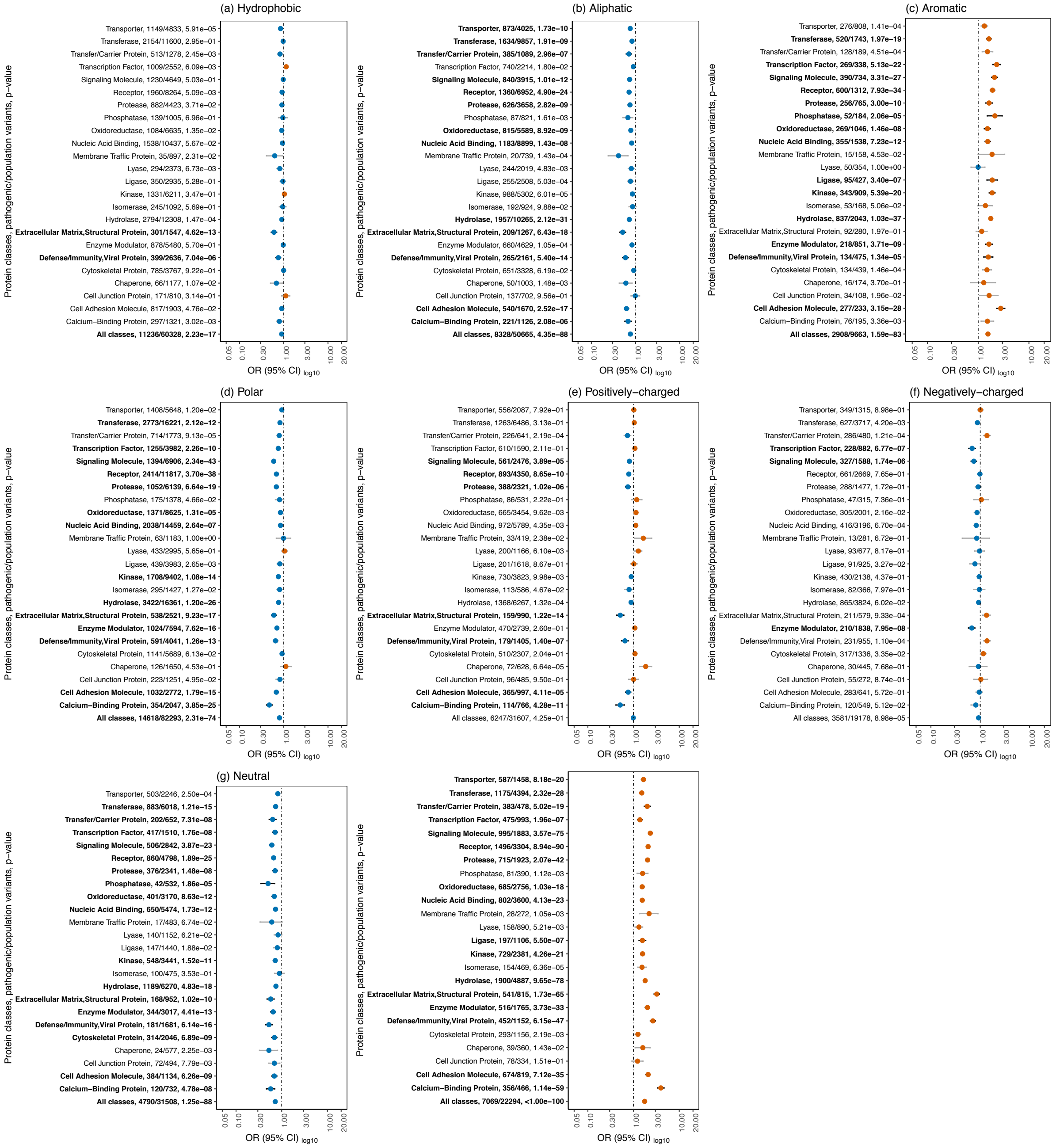


**Supplementary Fig. 5 | Protein-class-specific burden of pathogenic and population missense variants in the physicochemical property feature.** In each plot, the odds ratio (OR) equal to 1 indicates there is no association (Fisher's Exact test) between a variant type and a feature subtype, whereas OR > 1 (orange) and OR < 1 (blue) indicates that the pathogenic and population variants, respectively, are enriched in a particular feature subtype. The minimum and maximum values of OR set to 0.0 and 20.0, respectively, for visualization. The x-axis labels (protein class, feature counts with pathogenic and population variants, p-values) and the horizontal bars showing the confidence intervals of OR are bold-faced for significant p-values, lower than the cut-off (5.50e-05) after the Bonferroni correction.

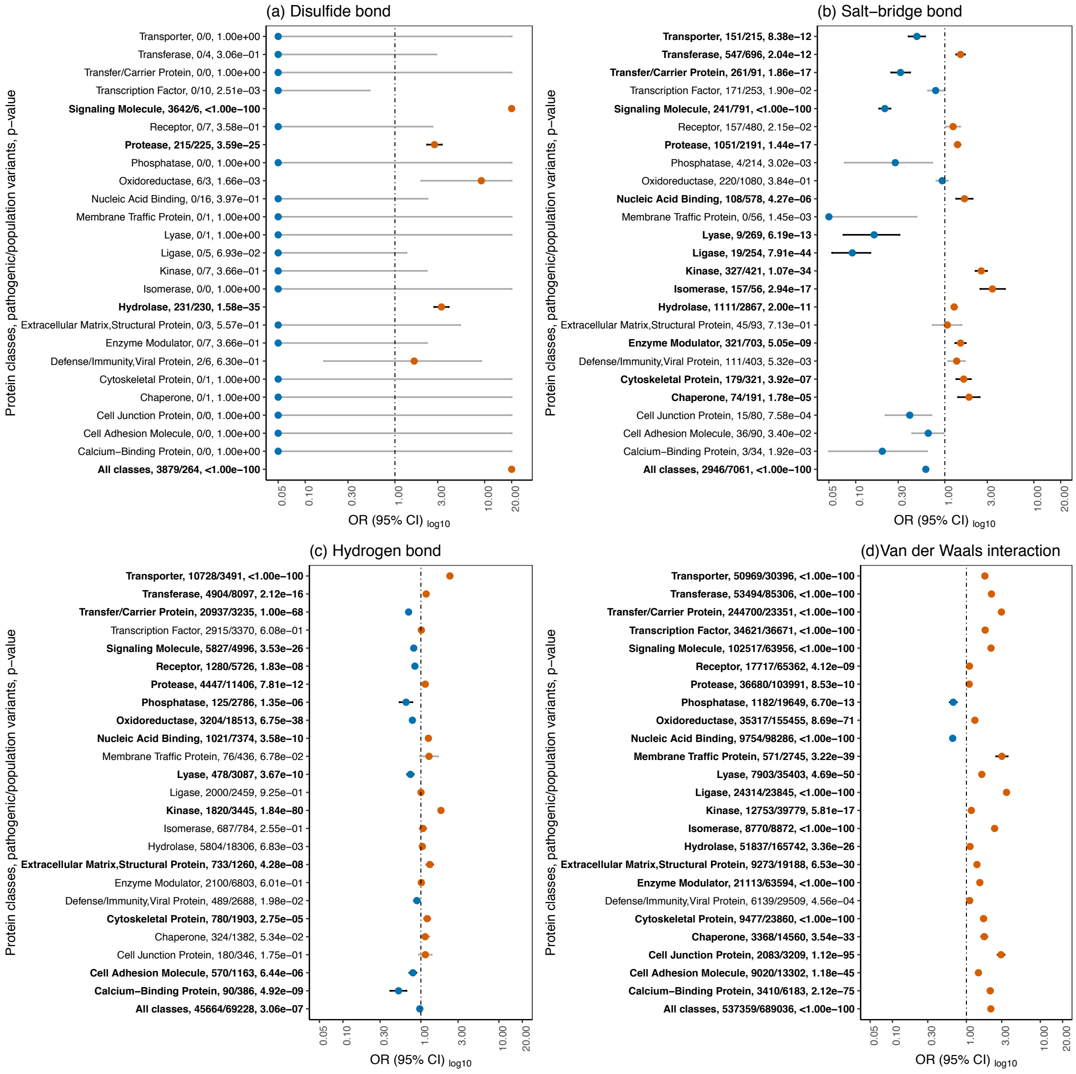


**Supplementary Fig. 6 | Protein-class-specific burden of pathogenic and population missense variants in the protein-protein interaction type feature.** In each plot, the odds ratio (OR) equal to 1 indicates there is no association (Fisher's Exact test) between a variant type and a feature subtype, whereas OR > 1 (orange) and OR < 1 (blue) indicates that the pathogenic and population variants, respectively, are enriched in a particular feature subtype. The minimum and maximum values of OR set to 0.0 and 20.0, respectively, for visualization. The x-axis labels (protein class, feature counts with pathogenic and population variants, p-values) and the horizontal bars showing the confidence intervals of OR are bold-faced for significant p-values, lower than the cut-off (5.50e-05) after the Bonferroni correction.

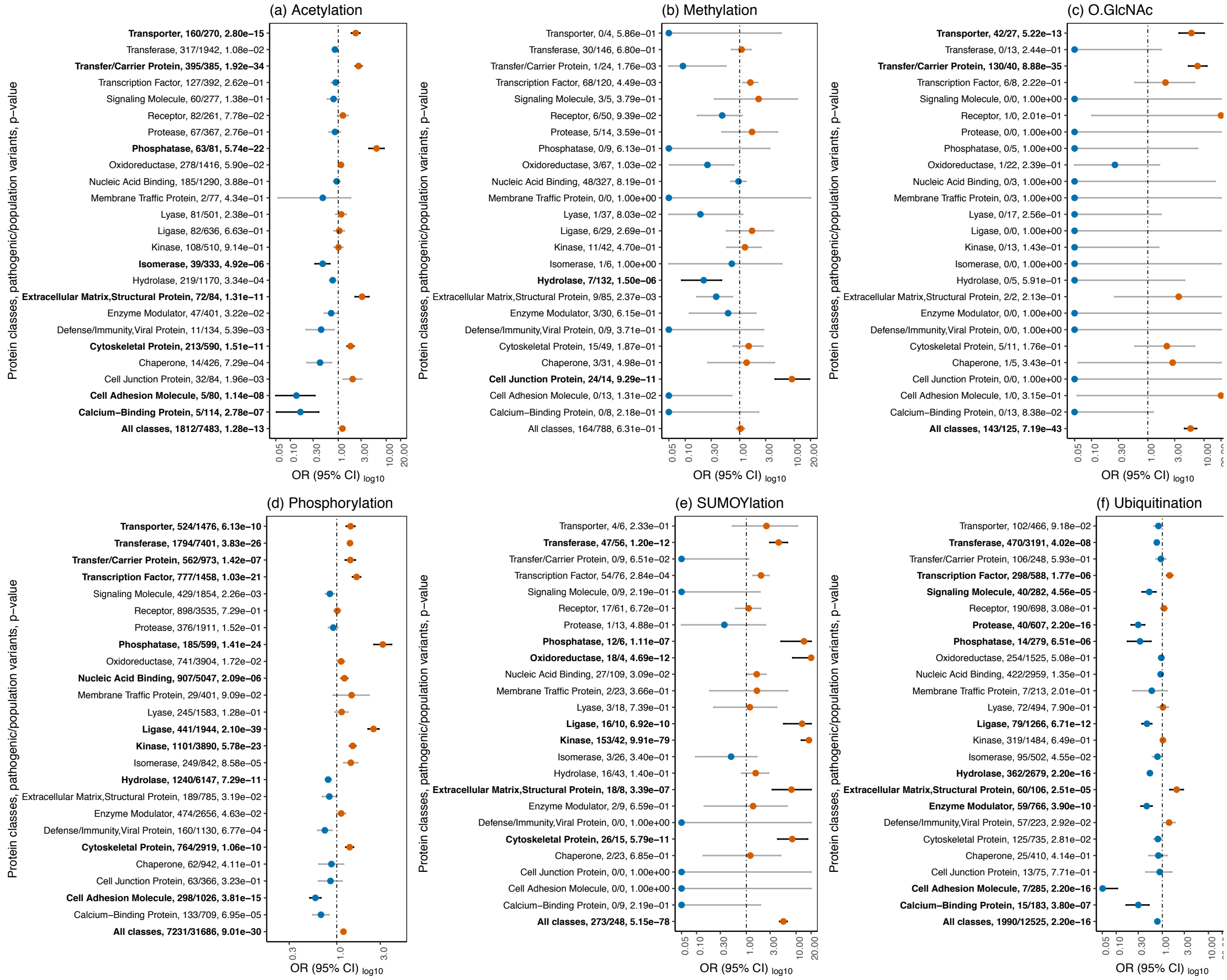


**Supplementary Fig. 7 | Protein-class-specific burden of pathogenic and population missense variants in the post-translational modification feature.** In each plot, the odds ratio (OR) equal to 1 indicates there is no association (Fisher's Exact test) between a variant type and a feature subtype, whereas OR > 1 (orange) and OR < 1 (blue) indicates that the pathogenic and population variants, respectively, are enriched in a particular feature subtype. The minimum and maximum values of OR set to 0.0 and 20.0, respectively, for visualization. The x-axis labels (protein class, feature counts with pathogenic and population variants, p-values) and the horizontal bars showing the confidence intervals of OR are bold-faced for significant p-values, lower than the cut-off (5.50e-05) after the Bonferroni correction.

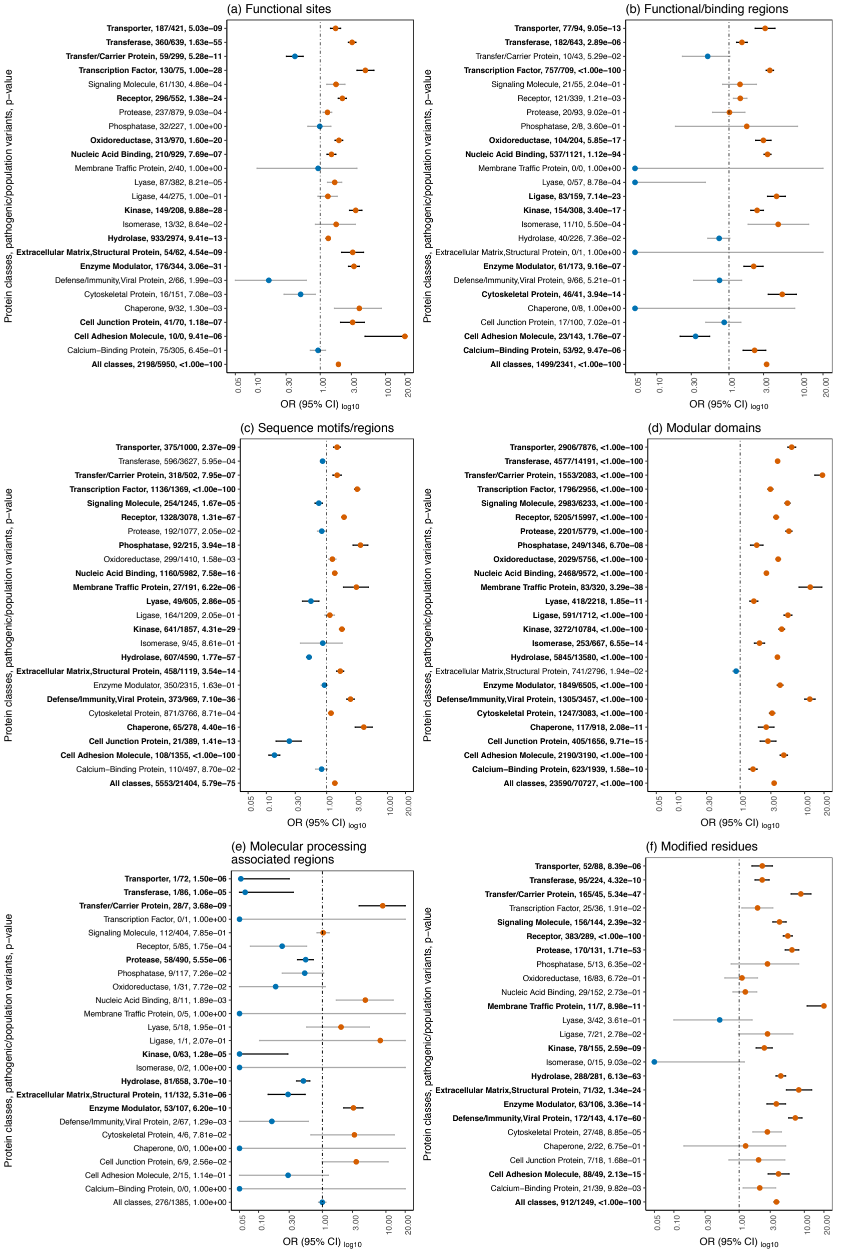


**Supplementary Fig. 8 | Protein-class-specific burden of pathogenic and population missense variants in the UniProt-based functional feature.** In each plot, the odds ratio (OR) equal to 1 indicates there is no association (Fisher's Exact test) between a variant type and a feature subtype, whereas OR > 1 (orange) and OR < 1 (blue) indicates that the pathogenic and population variants, respectively, are enriched in a particular feature subtype. The minimum and maximum values of OR set to 0.0 and 20.0, respectively, for visualization. The x-axis labels (protein class, feature counts with pathogenic and population variants, p-values) and the horizontal bars showing the confidence intervals of OR are bold-faced for significant p-values, lower than the cut-off (5.50e-05) after the Bonferroni correction.

(a)

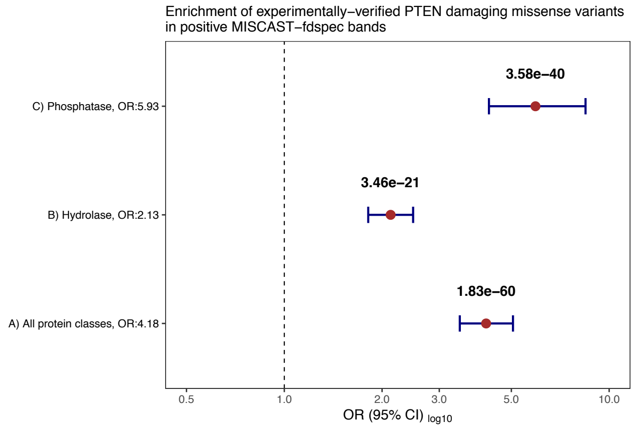


(b)

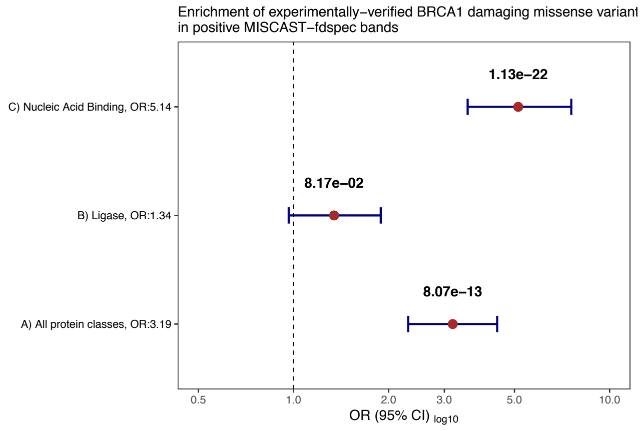


**Supplementary Fig. 9 | Validation of MISCAST-fdspec on experimentally-derived loss-of-function (or damaging) and neutral (or benign) missense variants, using high-throughput mutagenesis.** a, Burden of damaging missense variants (n = 876) in *PTEN* compared to neutral variants (n = 4,795) in positive MISCAST-fdspec bands (higher pathogenic variant-associated features), both along all-class and protein class-specific MISCAST-fdspec. *PTEN* is annotated as phosphatase and hydrolase, and the phosphatase protein class-specific features had the highest burden of damaging missense variants in *PTEN*. b, Burden of damaging variants (n = 248) in exon 13 of *BRCA1* compared to neutral variants (n = 1,381) in positive MISCAST-fdspec bands (higher pathogenic variant-associated features), both along all-class and protein class-specific MISCAST-fdspec. *BRCA1* is annotated as ligase and nucleic acid binding protein, and the nucleic acid binding protein class-specific features had the highest burden of damaging missense variants in *BRCA1*.

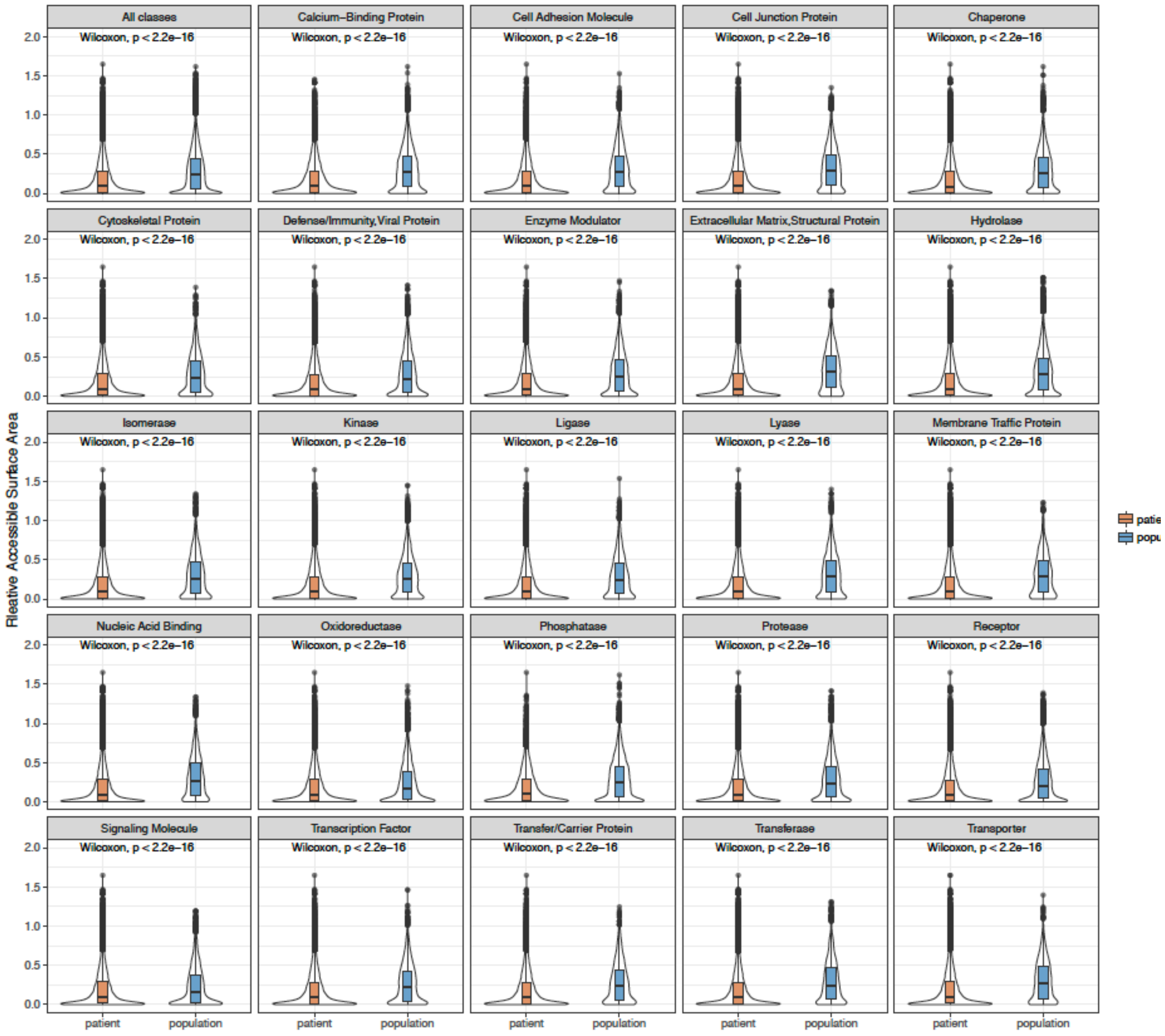


**Supplementary Fig. 10 | Distribution of relative accessible surface area (RSA) of amino acid residues on protein structure affected by population and pathogenic variants in 1,330 genes.** The significance of the Mann-Whitney rank sum test, showing the difference between the distributions of amino acids mutated by pathogenic and population variants, is shown in each plot. The residues mutated by pathogenic missense variants were found more in the interior of the protein, having at least 2-times lower median relative accessible surface area (RSA), than those mutated by population variants. Interestingly for majority of protein classes, the amino acid residues with pathogenic missense variants also accumulate higher number of outliers (high RSA values) which indicate the coexistence of pathogenic variants on protein surface, perturbing the essential interaction sites as well.

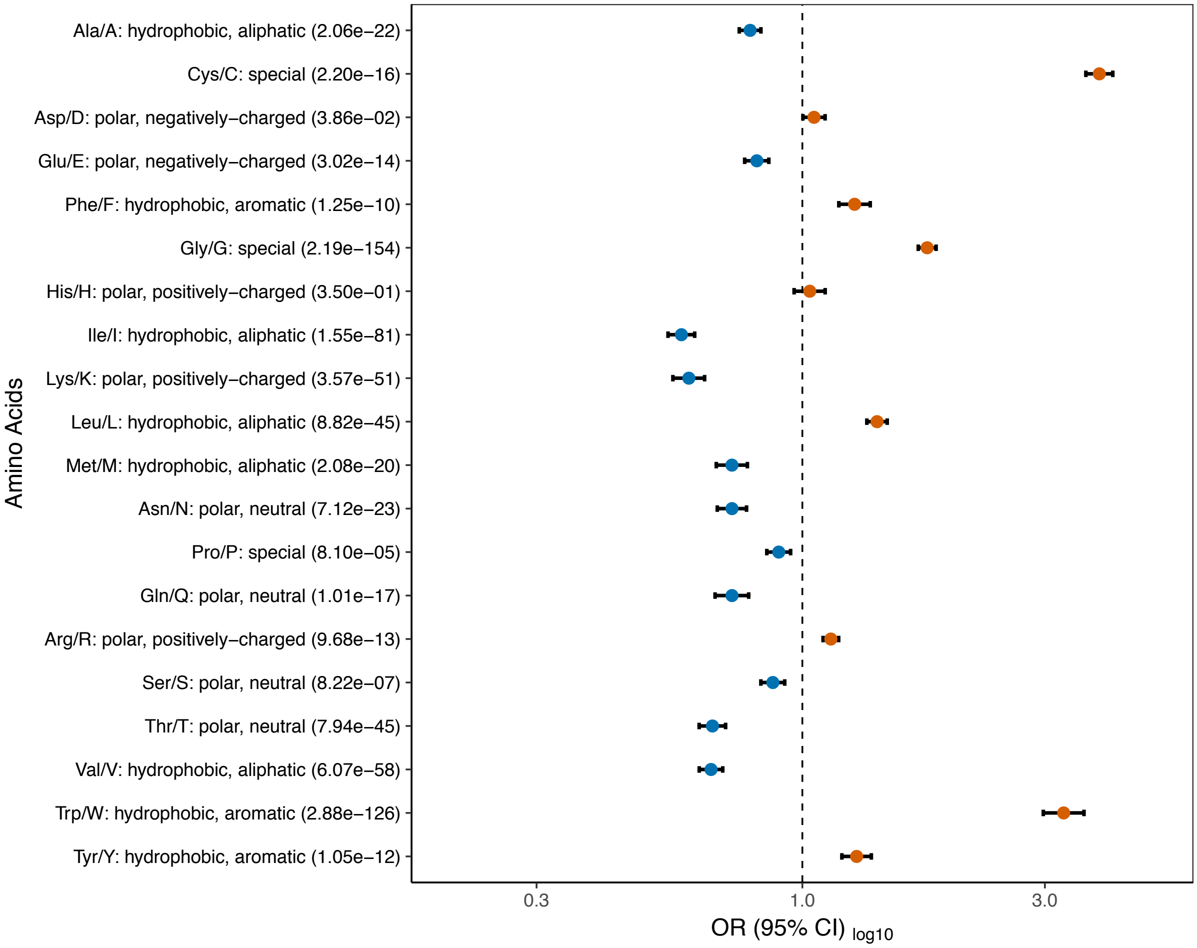


**Supplementary Fig. 11 | Association of pathogenic and population missense variants in 1,330 genes with 20 different amino acids.** In the plot, the odds ratio (OR) equal to one indicates there is no association (Fisher's Exact test) between a specific type of variant and a particular amino acid, whereas OR > 1 (orange) and OR < 1 (blue) indicates that the pathogenic variants are enriched (or depleted) in a particular type of amino acids and vice versa. The x-axis shows the OR and the 95% confidence interval, and the y-axis indicates the twenty amino acids, their physiochemical properties and p-values.

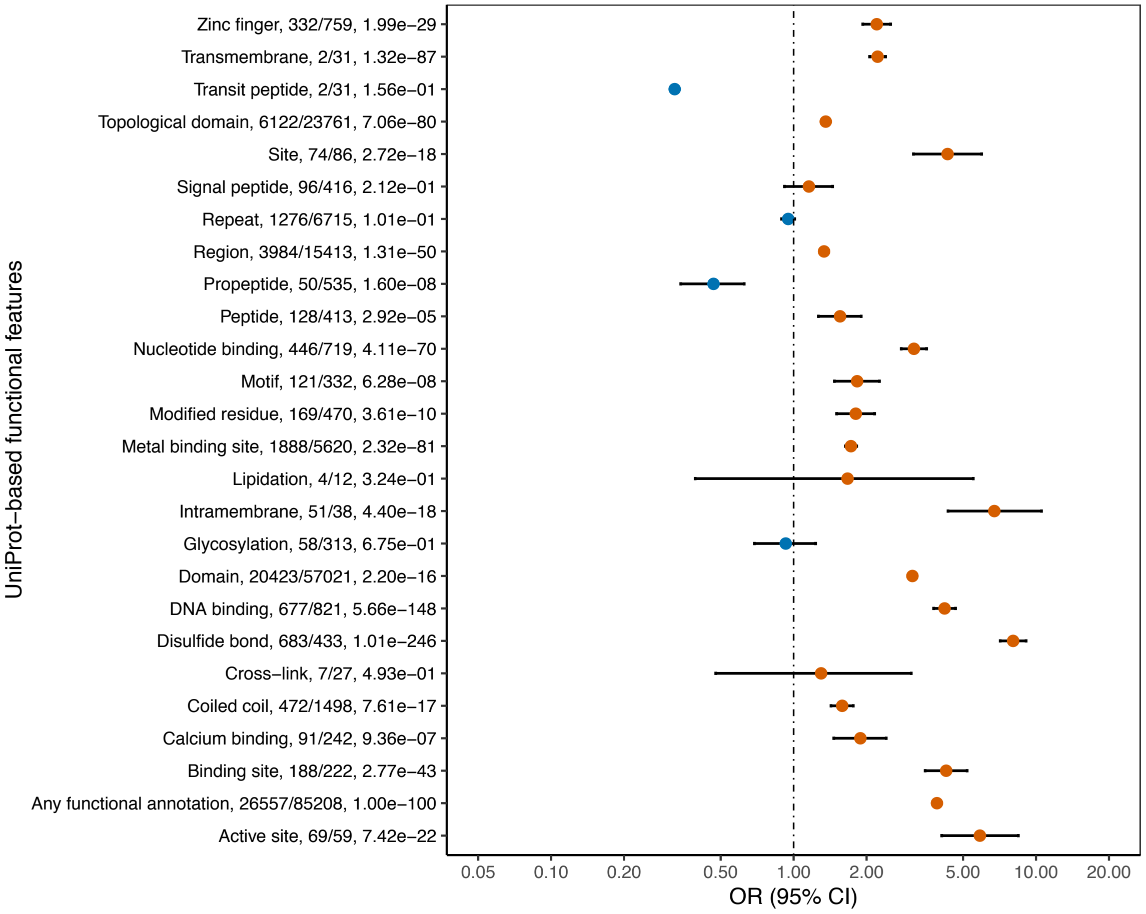


**Supplementary Fig. 12 | Association of pathogenic and population missense variants in 1,330 genes with 25 different UniProt-based functional features.** In the plot, the odds ratio (OR) equal to one indicates there is no association (Fisher's Exact test) between a specific type of variant and a particular amino acid, whereas OR > 1 (orange) and OR < 1 (blue) indicates that the pathogenic variants are enriched (or depleted) in a particular type of amino acids and vice versa. The x-axis shows the OR and the 95% confidence interval, and the y-axis indicates the twenty amino acids, their physiochemical properties and p-values. The output for any functional annotation shows the burden the pathogenic variants on any functional features versus no feature.
